# Modular split-trigger synergistic activation enables a programmable Cas13-12 system

**DOI:** 10.64898/2026.06.26.734747

**Authors:** Rui Sang, Avin Mohammadnejad Mavaneh, Ewa M. Goldys, Fei Deng

## Abstract

Achieving precise control of CRISPR-Cas *trans*-cleavage depends on understanding how nucleic acid activators engage Cas effectors, yet the fundamental principles of split-trigger activation of Cas12a remain unclear. Here, we uncover the mechanistic determinants that enable fragmented nucleic acids to collectively initiate Cas12a activity. We show that split triggers bearing external extensions fully support the R-loop formation, whereas internal extensions which disrupt the spacer complementarity abolish Csa12a activation. We further demonstrate that covalent linkage of split-trigger fragments prevents R-loop propagation, revealing that Cas12a’s activation strictly requires two physically independent split fragments. Together, these findings establish a synergistic split-trigger activation mechanism in which synergistic hybridization of two individually fragments nucleates and extends the Cas12a R-loop with high efficiency. Conceptually, this mechanism enables a Cas13-Cas12 architecture that transforms CRISPR diagnostics from a one-target one-Cas ribonucleoprotein (RNP) paradigm into a highly amplifying process in which a single target molecule activates numerous downstream Cas RNPs. Building on this principle, we show that the cleavage of a rationally designed linear DNA-RNA-DNA mediator by LbuCas13a generates optimally configured split triggers for Cas12a activation, thereby coupling RNA recognition to large-scale Cas12a activation without enzymatic preamplification. The resulting Split Trigger Activated Cas13-Cas12 System (STACS) achieves amplification-free detection down to 1 copy/µL within 15 minutes and maintains robust performance in complex biological (serum, saliva) and environmental (mud) matrices. This work establishes a generalizable strategy for engineering programmable CRISPR networks with high Cas RNP activation multiplicity for ultrasensitive molecular diagnostics.

## Introduction

Precise control of CRISPR-Cas *trans*-cleavage requires a detailed understanding of how nucleic acid activators engage Cas effectors (1–4). In CRISPR-based sensing, target recognition by a guide RNA initiates R-loop formation between the crRNA spacer and the complementary nucleic acid sequence, which the induces indiscriminate *trans*-cleavage activity of the Cas nuclease against single-stranded nucleic acid reporters (1,2,5). This target triggered signal amplification mechanism has extended the applications of CRISPR-Cas systems beyond gene editing enabling versatile molecular sensing and it underpins the sensitivity and specificity of CRISPR diagnostics (6–8). Consequently, the molecular rules governing Cas activation - rather than the downstream signal readout alone - represent a fundamental constraint in CRISPR-based nucleic acid detection.

Extensive efforts have been devoted to engineering nucleic acid activators to modulate Cas12a activity (9). Variations in activator length, nucleotide composition, and structural topology - including elongated activators, random 3′ extensions, dangling-end stabilization, DNA-RNA chimeric triggers, and sterically constrained architectures, have revealed that Cas12a activation is highly sensitive to trigger geometry and flexibility (10–14). These studies have significantly advanced the understanding of R-loop formation and *trans*-cleavage kinetics. However, the mechanistic insights obtained thus far are almost exclusively derived from contiguous activators. As a result, current design rules do not fully explain how two physically separate fragments of a nucleic acid activator, termed split triggers, can cooperate to activate Cas12a. In this arrangement, the two fragments bind to adjacent regions of the crRNA spacer and together promote activation, whereas either fragment alone exhibits minimal activity.

Recent studies provide initial evidence that split activators can modulate Cas12a activity under certain conditions. PAM-proximal helper strands have been shown to rescue Cas12a activation by replacing the DNA spacer region with RNA targets, enabling amplification-free detection of viral RNA and microRNAs (15). In parallel, synergistic activation has been observed when two short ssDNA fragments are simultaneously present, leading to enhanced *trans*-cleavage and improved mismatch discrimination (16). Similarly, combinations of two truncated ssDNA segments that individually exhibit minimal activity have been shown to synergistically restore Cas12a trans-cleavage, suggesting that complementary engagement of different regions of the crRNA spacer can support Cas12a activation (17). Nevertheless, these reports primarily describe phenomenological activation behavior and do not establish generalizable molecular rules governing fragment orientation, extension position, physical linkage, or R-loop propagation. As a result, the principles by which split triggers nucleate and extend the Cas12a R-loop remain undefined.

Here, we systematically define the molecular rules governing Cas12a activation by split nucleic acid triggers. We demonstrate that split triggers bearing external nucleotide extensions fully support R-loop formation, whereas internal extensions disrupt spacer complementarity and abolish activation. We further establish that Cas12a activation strictly requires two physically independent fragments, as covalent linkage prevents productive R-loop propagation. By systematically mapping fragment length effects, we reveal a division of labour in which one fragment nucleates R-loop initiation while the complementary fragment promotes R-loop extension and stabilization. Together, these findings establish a mechanistic framework for synergistic split-trigger mediated Cas12a activation.

Furthermore, we show that, as a direct consequence of these mechanistic constraints, the cleavage of a specially designed linear DNA-RNA-DNA mediator by LbuCas13a generates split fragments that are optimally configured for Cas12a activation. Leveraging this effect, we construct a Split Trigger Activated Cas13-Cas12 System (STACS), which transforms RNA recognition into sequential CRISPR amplification without enzymatic preamplification. Unlike conventional amplification-assisted CRISPR diagnostics(15,18), STACS achieves detection without target preamplification down to 1 copy µL□¹ of SARS-CoV-2 RNA within 15 minutes after sample preparation, with robust performance across complex biological and environmental matrices, including serum, saliva, and mud. Importantly, the linear mediator is prepared using standard commercial methods and is readily obtainable from many suppliers. Conceptually, this design transforms CRISPR diagnostics from a conventional one-target one-Cas ribonucleoprotein (RNP) paradigm into a sequential activation architecture in which a single target molecule triggers the activation of numerous downstream Cas RNPs. This work bridges fundamental CRISPR activation biology with practical diagnostic engineering and provides a mechanistic foundation for designing programmable CRISPR systems with sequential signal transfer.

## Materials and Methods

All nucleic acid sequences listed in Table 1 were purchased from Gencefe Biotechnology Co. Ltd (Jiangsu, China). LbuCas13a protein (Z03742-1) was purchased from GenScript Biotech (Nanjing, China). LbCas12a protein (M0653T) and rCutSmart buffer (B6004S) were purchased from NEB Biotechnology Co. (Ipswich, MA UK). DNase/RNase Free Water (10977015), Proteinase K (25530049) were purchased from Thermo Fisher Scientific (Waltham, MA US). Human serum (H4522) was purchased from Sigma-Aldrich (St. Louis, Missouri US). Milenia HybriDetect LFA kit (MGHD 1) was purchased from Milenia Biotec Gmbh (Giessen, Germany). Inactivated SARS-CoV-2 reference material (NMIA NA050-NA055, B200921) were provided by Australian Government National Measurement Institute.

**Table 1.**
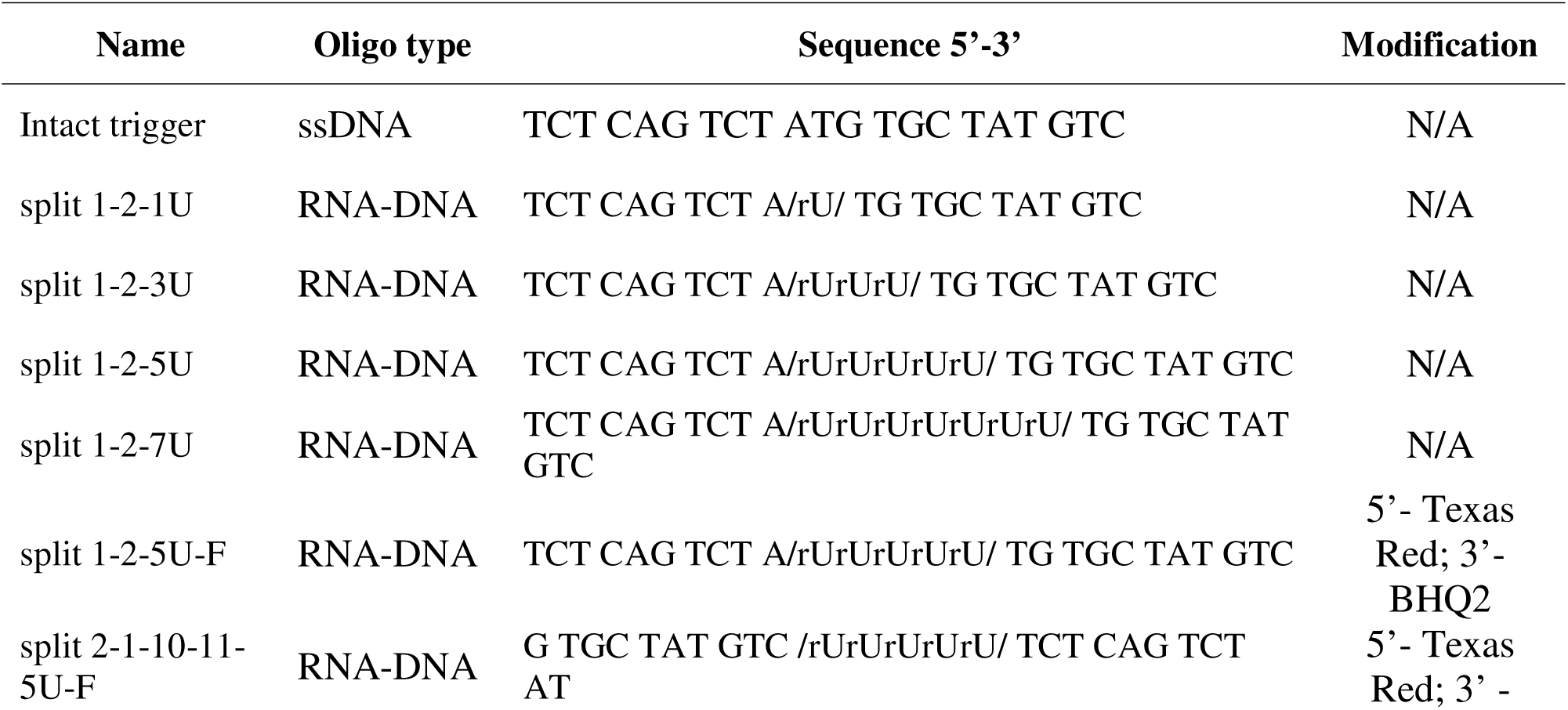
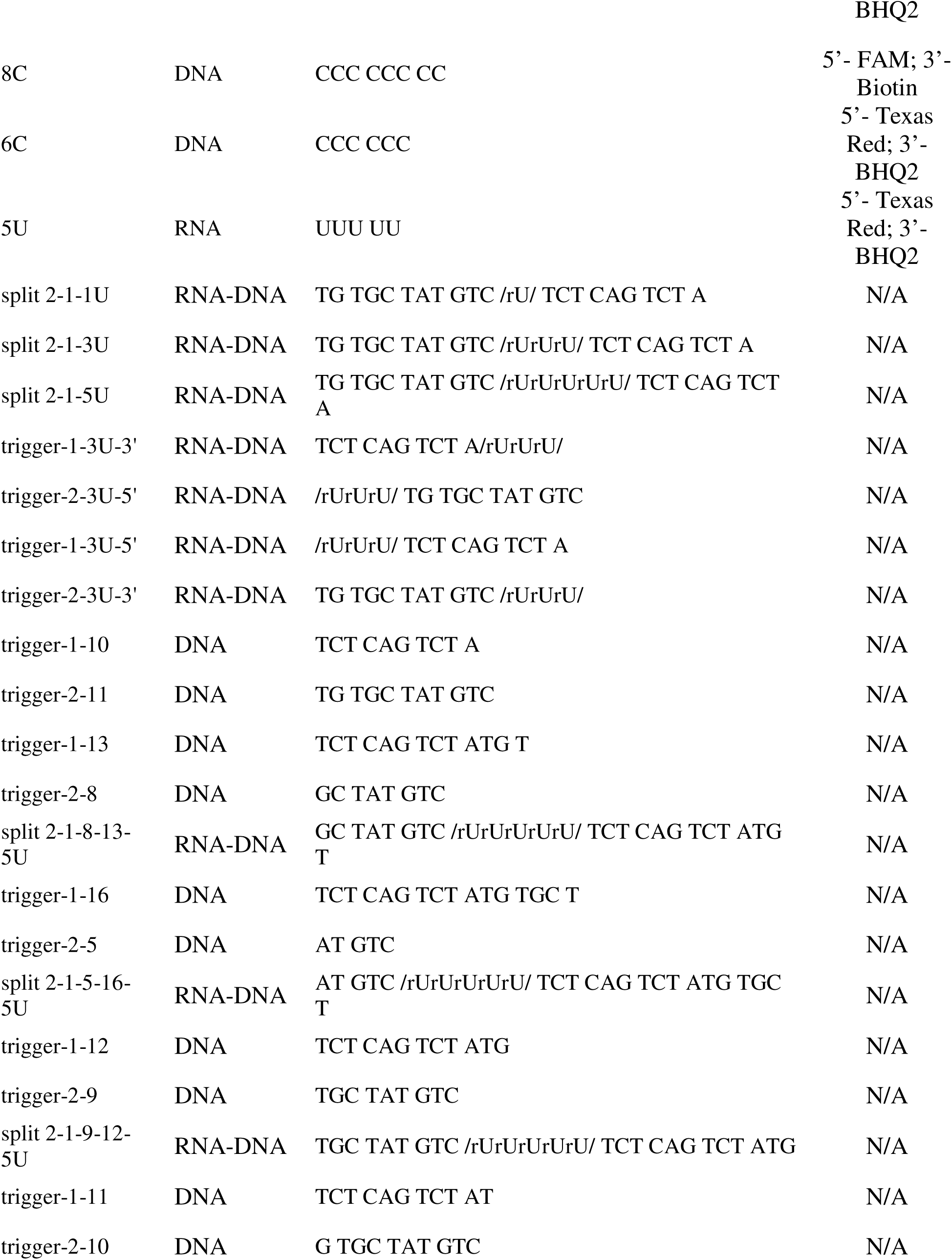
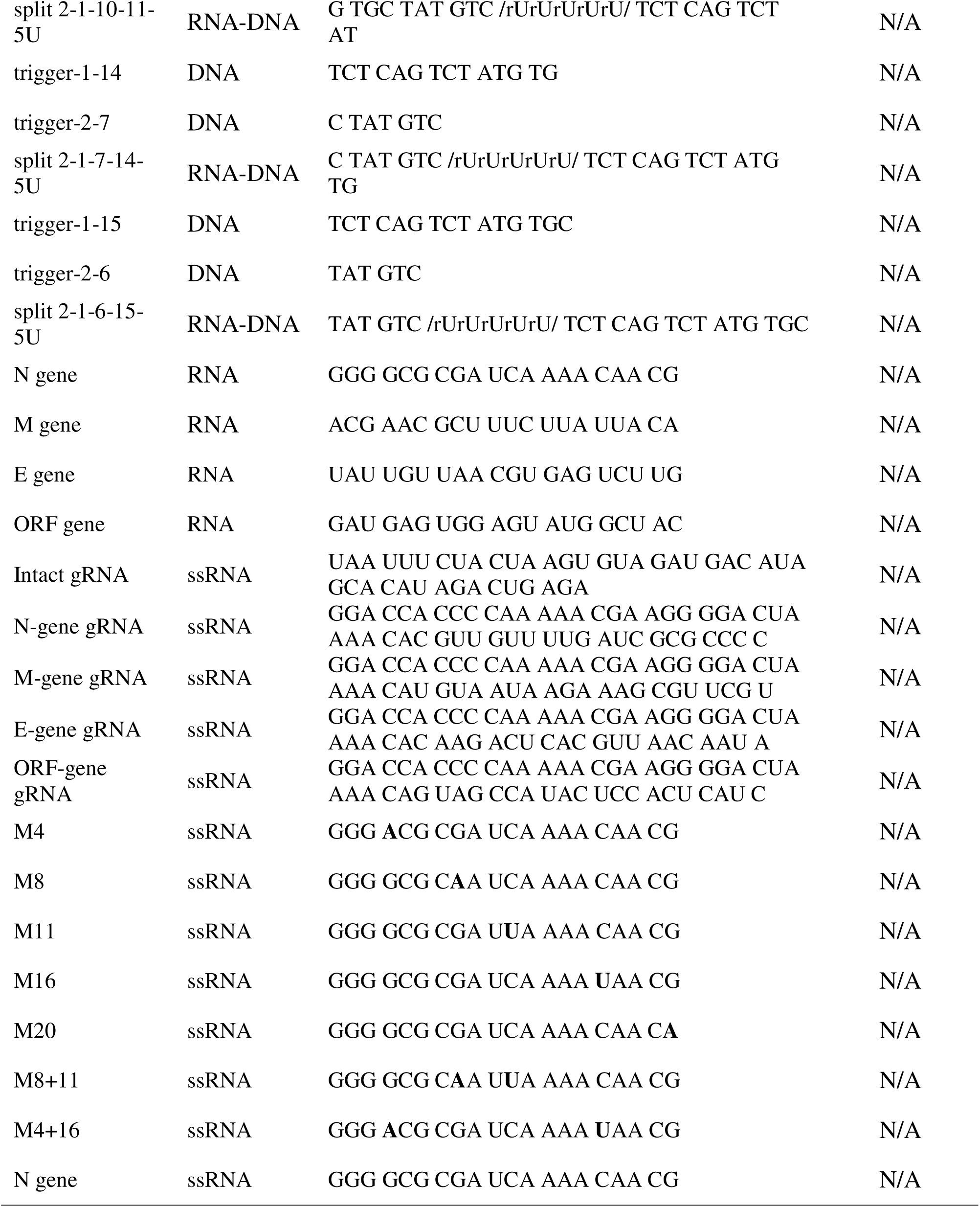
DNA and RNA oligos are utilized in this study.

### Preparation of the standard CRISPR reaction solution

The standard CRISPR reaction solution of LbCas12a and LbuCas13a were prepared as follows:

#### LbCas12a

1 μL of 20 μM of LbCas12a endonuclease and 1 μL of 20 μM of Intact trigger gRNA were mixed at 1 mL 1× rCutSmart buffer, followed by addition of 5 μL of 100 μM of 6C reporter. The prepared standard CRISPR/Cas12a reaction mixture was preincubated for 30 min at 37□and then stored at 4□for future use.

#### LbuCas13a

1 μL of 20 μM of LbuCas13a endonuclease and 1 μL of 20 μM of LbuCas13a SARS-CoV-2 gRNA were mixed at 1 mL 1× rCutSmart buffer, followed by addition of 5 μL of 100 μM of split mediator or 5U reporter. The prepared standard CRISPR/Cas13a reaction mixture was preincubated for 30 min at 37□and then stored at 4□for future use.

### Investigation of trigger mechanism of split mediator for Lbcas12a RNP

To investigate different split mediators’ activation mechanism for Lbcas12a, 2μL of 1μM of trigger (split 1, split 2, split 1 & split 2 and split mediators) was added into 98 μL of standard CRISPR/ LbCas12a reaction solution (20nM of LbCas12a protein, 20nM of gRNA, 500nM of reporter, and 1mL of rCutSmart buffer). The reaction was carried out at 37□, and the fluorescence intensity at Ex/Em of 570/615 nm was determined by using a plate reader (iD5 Spectramax, Molecular Devices, USA).

#### Multiple gRNA reaction buffer

The CRISPR reaction solution of LbCas12a and LbuCas13a for multiple gRNA reaction were prepared as follows:

#### LbuCas13a

4 μL of 20 μM of LbuCas13a endonuclease and 4 μL of 20 μM of LbuCas13a SARS-CoV-2 gRNA (specific to N gene, M gene, E gene or ORF gene) were mixed at 1 mL 1× rCutSmart buffer, followed by preincubation for 30 min at 37□. Then, these four different RNP solutions were mixed and 6 μL of 10 μM of split mediator were added and stored at 4□for future use.

#### 10×LbCas12a

40 μL of 20 μM of LbCas12a endonuclease and 40 μL of 20 μM of Intact trigger gRNA were mixed at 1 mL 1× rCutSmart buffer, followed by addition of 1.66 μL of 100 μM of 6C/8C reporter. The prepared standard CRISPR/Cas12a reaction mixture was preincubated for 30 min at 37□and then stored at 4□for future use.

### Detection of synthetic SARS-CoV-2 genomic RNA

To investigate the LOD of STACS for synthetic SARS-CoV-2 genomic RNA detection, 10 μL of 10,000 copies/mL, 1,000 copies/mL, 100 copies/mL of synthetic SARS-CoV-2 genomic RNA was added into 90 μL of standard multiple gRNA CRISPR/ LbuCas13a reaction solution for activating *trans*-cleavage of LbuCas13a. The reaction mixture was incubated at 37□for 30 min, followed by adding 10 μL of 10 times standard multiple gRNA CRISPR/ LbCas12a reaction solution. The fluorescence intensity at Ex/Em of 570/615 nm was determined by using a plate reader (iD5 Spectramax, Molecular Devices, USA) at 37□.

### Detection of synthetic SARS-CoV-2 genomic RNA in human serum

Proteinase K treatment: 40 μL of human serum were mixed with 20 μL of proteinase K (20mg/ml). The reaction solutions were incubated at 55□for 30 min and then 95 for 15 min, followed by cooling down to room temperature.

To investigate the detection ability of STACS for synthetic SARS-CoV-2 genomic RNA in 10% human serum, 10 μL of 10,000 copies/mL, 1,000 copies/mL, 100 copies/mL of synthetic SARS-CoV-2 genomic RNA in DPBS with 10% human serum was added into 90 μL of standard multiple gRNA-CRISPR/LbuCas13a reaction solution for activating *trans*-cleavage of LbuCas13a. The reaction mixture was incubated at 37□for 30 min, followed by adding 10 μL of 10 times standard multiple gRNA-CRISPR/LbCas12a reaction solution. The fluorescence intensity at Ex/Em of 570/615 nm was determined by using a plate reader (iD5 Spectramax, Molecular Devices, USA) at 37□.

All human serum experiments were approved by the UNSW Ethics Committee (UNSW HC210160).

### Detection of synthetic SARS-CoV-2 genomic RNA in environmental samples

Sample preparation was carried out as follows. Environmental mud samples were diluted 10 times in water and then filtered by 0.22 μm syringe filter and stored at 4□until further use.

To investigate the detection ability of STACS for synthetic SARS-CoV-2 genomic RNA in environmental samples, 10 μL of 10,000 copies/mL, 1,000 copies/mL, 100 copies/mL of synthetic SARS-CoV-2 genomic RNA in 1 × rCutSmart with 5% mud was added into 90 μL of standard multiple gRNA-CRISPR/LbuCas13a reaction solution for activating *trans*-cleavage of LbuCas13a. The reaction mixture was incubated at 37□for 30 min, followed by adding 10 μL of 10 times standard multiple gRNA-CRISPR/LbCas12a reaction solution. The fluorescence intensity at Ex/Em of 570/615 nm was determined by using a plate reader (iD5 Spectramax, Molecular Devices, USA) at 37□.

### Detection of inactivated SARS-CoV-2 reference RNA

To investigate the detection ability of STACS for inactivated SARS-CoV-2 reference material, 10 μL of 100,610 copies/mL, 10,750 copies/mL, 1,120 copies/mL or 360 copies/mL of inactivated SARS-CoV-2 reference material was added into 90 μL of standard multiple gRNA CRISPR/LbuCas13a reaction solution. The reaction mixture was incubated at 37□for 30 min, followed by adding 10 μL of 10 times standard multiple gRNA-CRISPR/LbCas12a reaction solution. The fluorescence intensity at Ex/Em of 570/615 nm was determined by using a plate reader (iD5 Spectramax, Molecular Devices, USA) at 37□.

### Detection of inactivated SARS-CoV-2 reference material in human saliva

Proteinase K treatment of SARS-CoV-2-spiked human saliva was performed as follows. Briefly, 4 μL of proteinase K (20 mg/mL) was mixed with 32 μL of undiluted human saliva and 4 μL of certified SARS-CoV-2 reference material containing inactivated virus (100,610 copies/mL; 42,500 copies/mL; 10,750 copies/mL; 3,210 copies/mL), or DPBS (as negative control). The mixture was incubated at 65□for 15 min, followed by 95 °C for 5 min and then cooled to 4 °C until further use.

To evaluate STACS detection in a biologically complex matrix following sample processing, 10 μL of the processed saliva sample containing SARS-CoV-2 reference material at 10,061 copies/mL, 4,250 copies/mL, 1,075 copies/mL or 321 copies/mL was added into 90 μL of standard multiple gRNA CRISPR/ LbuCas13a reaction solution. The reaction mixture was incubated at 37□for 30 min, followed by adding 10 μL of 10 times standard multiple gRNA-CRISPR/ LbCas12a reaction solution. The fluorescence intensity at Ex/Em of 570/615 nm was determined by using a plate reader (iD5 Spectramax, Molecular Devices, USA) at 37□.

All human saliva experiments were approved by the UNSW Ethics Committee (UNSW HC 200568).

### Lateral flow assays (LFA)

To investigate the detection ability of STACS for SARS-CoV-2 Standards RNA via LFA, 10 μL of synthetic SARS-CoV-2 genomic RNA or inactivated SARS-CoV-2 reference material was added into 90 μL of standard multiple gRNA-CRISPR/LbuCas13a reaction solution for activating *trans*-cleavage of LbuCas13a. The reaction mixture was incubated at 37□for 30 min, followed by adding 10 μL of 10 times standard multiple gRNA-CRISPR/ LbCas12a reaction solution (using 8C as reporter). Then, 6 μL of reaction solution were mixed with 94 μL of Milenia running buffer at 2 min, 5 min, 7 min, 15 min, 30 min, followed by adding Milenia HybriDetect 1 strip into the tube and allowed to run for around 3 minutes. Lastly, strips were read on a lateral Flow Reader (AX-2X-S, Axxin Pty Ltd, Australia).

## Results

### 1. External-Extension split triggers efficiently activate LbCas12a

We first evaluated whether fragmented activators could support Cas12a *trans*-cleavage. The two unmodified split fragments of a fully competent activator (trigger-1 and trigger-2) restored full activation when supplied together, yielding fluorescence kinetics equivalent to the intact activator, whereas either fragment alone showed minimal activity (Fig.1A and Fig. S1). These results confirm that Cas12a efficiently recognises a bipartite activator, provided that the complementary region required for R-loop formation remains intact.

**Figure 1.**
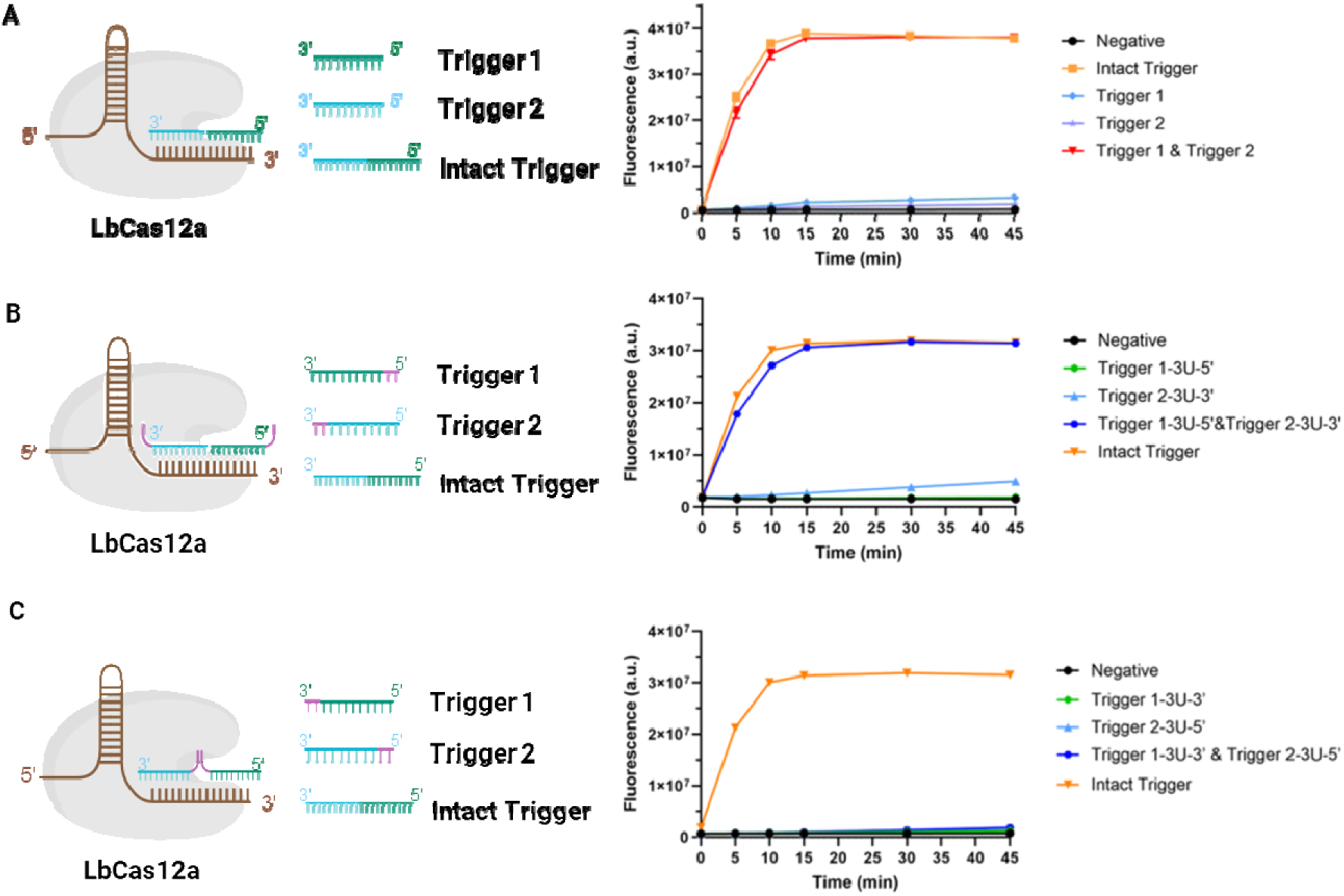
Split triggers with external extensions are able to fully activate LbCas12a. (A) Split triggers are able to fully activate LbCas12a; (B) Split triggers with external extensions are able to fully activate LbCas12a; (C) Split triggers with internal extensions are not able to activate LbCas12a. n=3, error bars represent mean ± SD, a.u = arbitrary units.

To determine how nucleotide extensions influence this process, we compared split triggers carrying either external or internal poly-U additions. External extensions positioned at the 5′ end of trigger-1 or the 3′ end of trigger-2 preserved robust activation: individually inactive, the two extended fragments together triggered Cas12a with efficiency comparable to the intact activator (Fig.1B). In contrast, internal U-extensions inserted between the two complementary halves completely abolished activation, even when both fragments were supplied (Fig.1C). This loss of activity indicates that internal insertions obstruct productive pairing with the crRNA spacer and disrupt R-loop propagation.

Together, the structural schematics and kinetic data in Fig. 1 establish a clear mechanistic rule: Cas12a tolerates nucleotide extensions only when they lie outside the crRNA-target pairing region, whereas internal extensions disrupt R-loop formation and abolish activation. This positional requirement provides a straightforward design principle for constructing functional split triggers.

### 2. Linked split triggers fail to fully activate LbCas12a

Having established that Cas12a tolerates *external* extensions on independent split fragments of trigger sequences, we next investigated whether these fragments could be covalently linked without compromising activation. To test this, we introduced poly-U linkers of varying lengths between trigger-1 and trigger-2 in two orientations (1→2 and 2→1), generating single-stranded constructs that retain full sequence complementarity but eliminate fragment independence (Fig. 2A-B).

**Figure 2.**
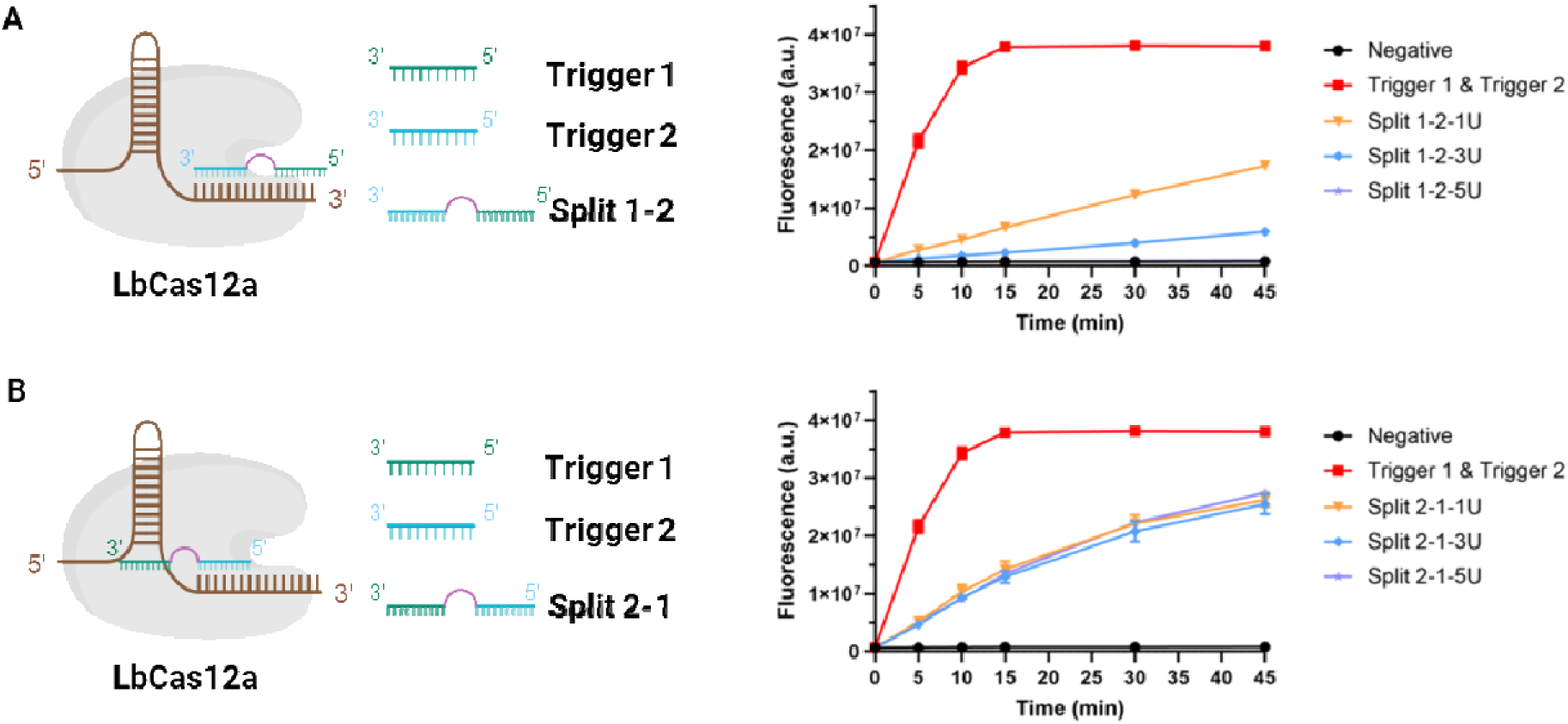
Linked split triggers are not able to fully activate LbCas12a. (A) The activation assessment of Split 1-2 with different U linker length for LbCas12a; (B) The activation assessment of Split 2-1 with different U linker length for LbCas12a. n=3, error bars represent mean ± SD, a.u = arbitrary units.

Across all linker lengths examined, linked split triggers displayed markedly reduced activation compared with the unlinked trigger-1 + trigger-2 pair. When trigger-1 was positioned at the 5′ end (Split 1-2 constructs), increasing the U-linker length from 1-U to 5-U leads to lower activation efficiency, and none approached the activation level achieved by the two independent fragments (Fig. 2A). When the orientation was reversed (Split 2-1 constructs), all linked versions generated only partial activation despite increasing linker flexibility (Fig. 2B).

These results reveal a mechanistic constraint distinct from that observed for external extensions. Whereas Cas12a tolerates nucleotide additions outside the pairing region, it does not tolerate physical linkage between trigger fragments. We infer that covalent linkage restricts the conformational freedom required for sequential binding of the two halves and prevents the stepwise R-loop propagation observed with independent fragments.

Together, these findings establish that Cas12a activation requires two physically separate oligonucleotides, which bind the crRNA-guided complex in a coordinated but structurally unconstrained manner.

### 3. Defining the length requirements for functional split-trigger activation of LbCas12a

To define the length parameters governing efficient synergistic activation of LbCas12a, we systematically varied the lengths of trigger-1 and trigger-2 while maintaining their relative positions within the target sequence (Fig. 3A-H). For each trigger pair, Cas12a activation by the independent trigger-1 and trigger-2 fragments was compared with that produced by the corresponding linked split mediator, allowing direct assessment of synergy and background activation.

**Figure 3.**
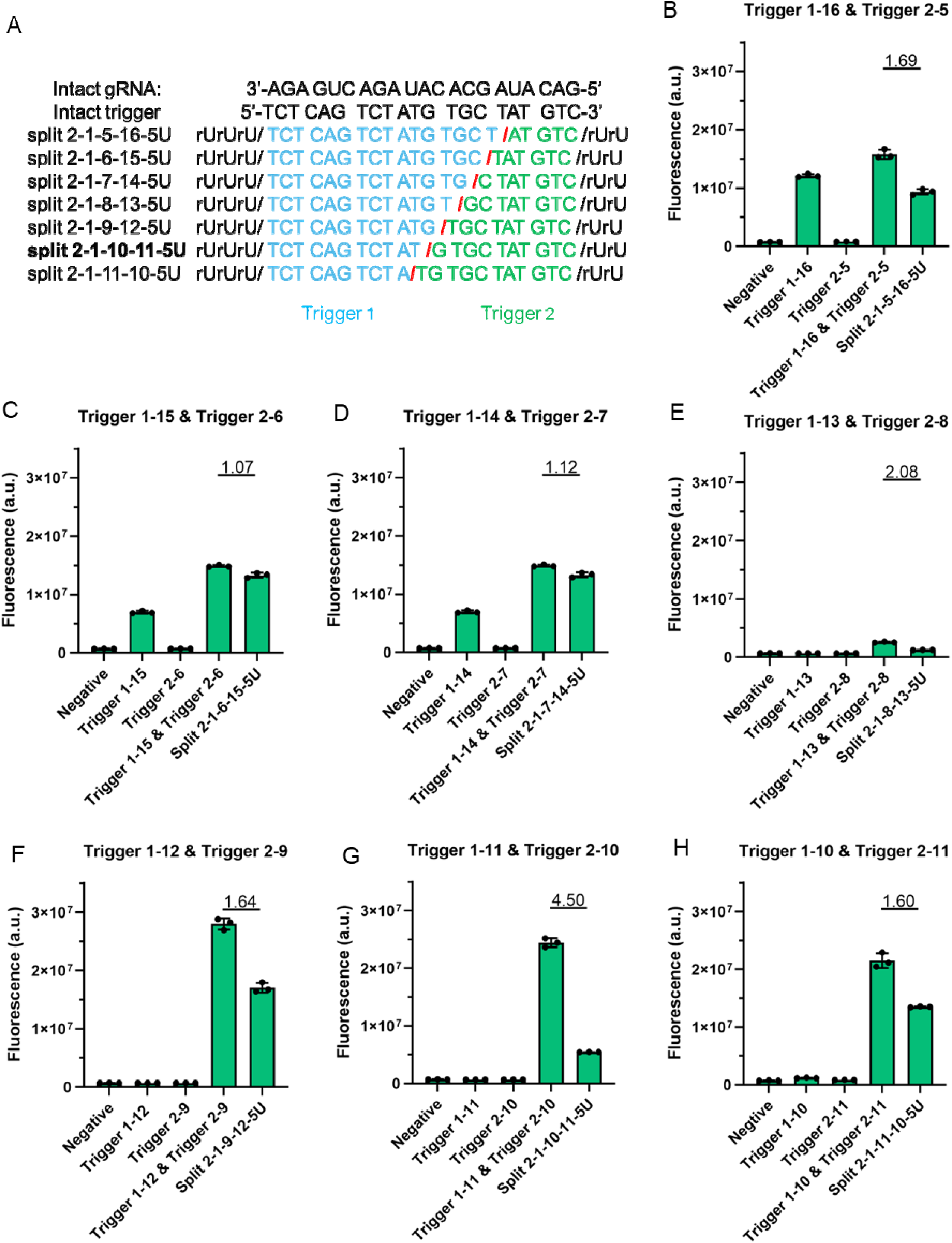
Defining the optimal length of trigger 1 and trigger 2. (A). Sequences of linear mediators with different lengths; (B-H). The ability of trigger 1 and trigger 2 with different lengths to activate LbCas12a. n=3, error bars represent mean ± SD, * P ≤ 0.05, ** P ≤ 0.01, *** P ≤ 0.001, **** P ≤ 0.0001, ns = non-significant, a.u = arbitrary units.

When trigger-1 was long (16-14 nt), trigger-1 alone was able to partially activate LbCas12a, and combining trigger-1 with trigger-2 resulted in only modest signal enhancement (Fig. 3B-D). In these configurations, the signal ratios between the paired triggers and the linked split mediators remained close to unity, indicating limited synergistic gain. This suggests that long trigger-1 fragments can independently nucleate R-loop formation, thereby reducing the relative contribution of synergistic activation.

As the length of trigger-1 was reduced to 13-12 nt, its standalone activation ability sharply decreased, and Cas12a activity became increasingly dependent on the presence of trigger-2 (Fig. 3E-F). Under these conditions, the paired trigger-1/trigger-2 combinations generated higher signal ratios, consistent with a transition from single-fragment activation to split-trigger-mediated R-loop assembly.

Maximal synergistic activation was observed when trigger-1 and trigger-2 were 11 nt and 10 nt in length, respectively (Fig. 3G). This configuration produced the highest fluorescence signal ratio (4.50), indicating optimal balance between insufficient standalone activation and efficient R-loop formation supported by the two split fragments. Further decrease of trigger-1 length to 10 nt reduced the signal ratio (Fig. 3H), suggesting that excessively short fragments compromise the initial R-loop nucleation even in the presence of the complementary trigger fragment.

Collectively, these results define a clear length-dependent division of tasks between the split fragments: trigger-1 primarily governs the R-loop nucleation, while trigger-2 supports the R-loop extension and stabilization. An optimal window exists in which trigger-1 is sufficiently short to suppress background activation but long enough to initiate synergistic binding with trigger-2. The 11 nt + 10 nt split configuration best satisfies these constraints and was therefore selected as the optimal split-trigger design for subsequent STACS development.

### 4. Performance evaluation of STACS

In STACS (Fig. 4A), target RNA is first recognized by sequence-programmed LbuCas13a, triggering robust collateral RNase activity. Each activated Cas13a RNP cleaves numerous linear DNA-RNA-DNA mediators, converting a single RNA recognition event into the generation of a large pool of physically independent split DNA fragments with external extensions. These cleavage products act as synergistic split triggers for LbCas12a, in which one fragment nucleates R-loop initiation while the complementary fragment promotes R-loop extension and stabilization, together activating otherwise inactive Cas12a RNPs. Consequently, numerous new Cas12a RNPs are activated downstream of a single target RNA molecule, transforming the conventional one-target one-Cas RNP paradigm into a sequential process with high Cas RNP activation multiplicity. The massively amplified Cas12a trans-cleavage activity then produces strong fluorescence or lateral-flow signals through reporter cleavage, enabling amplification-free, low-background, and ultrasensitive nucleic acid detection.

**Figure 4.**
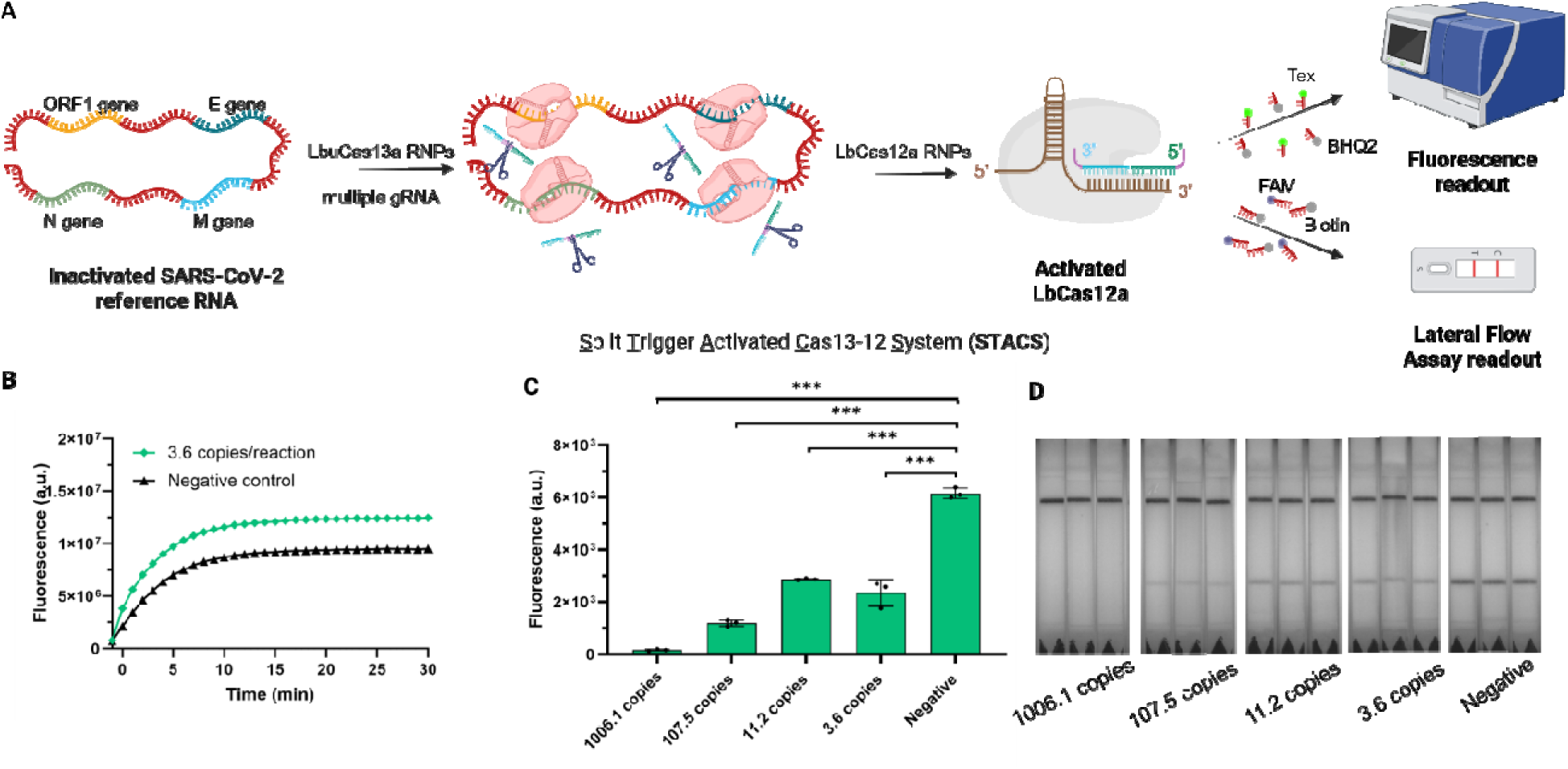
Performance evaluation of STACS using a multiple-gRNA strategy. (A) Schematic illustration of STACS employing four LbuCas13a gRNAs targeting the N, M, E, and ORF gene of inactivated SARS-CoV-2 reference material. (B) Fluorescence kinetics of STACS for detection of 3.6 copies of inactivated SARS-CoV-2 reference material. (C) The detection range of STACS for inactivated SARS-CoV-2 reference material using LFA. (D). The corresponding images of figure C. n=3, error bars represent mean ± SD, * P ≤ 0.05, ** P ≤ 0.01, *** P ≤ 0.001, **** P ≤ 0.0001, ns = non-significant, a.u = arbitrary units.

Based on this architecture, the performance of STACS for SARS-CoV-2 RNA detection was evaluated using a multiple-gRNA recognition strategy (Fig. 4A). The *trans*-cleavage ability of LbuCas13a on linear mediator was demonstrated via both agarose gel electrophoresis analysis (Fig. S2) and fluorescent readout (Fig. S3) and the optimum buffer for STACS was found to be rCutSmart (Fig. S4). Four LbuCas13a gRNAs targeting the N, M, E, and ORF regions of the viral genome were combined to enhance target engagement and maximize Cas13a activation, thereby increasing the probability of productive downstream Cas12a activation (Fig. S5). In addition, the incubation time of STACS was optimized to be close to 15min (Fig. S6).

Using this configuration, STACS enabled rapid and sensitive detection of reference SARS-CoV-2 RNA targets by using plate reader (Fig. S7). When 3.6 copies of inactivated SARS-CoV-2 reference material were present per reaction, a clear and time-dependent fluorescence increase was observed compared with the negative control (Fig. 4B), confirming a successful sequential activation at near single-copy levels without enzymatic preamplification.

To demonstrate the relevance of STAC in a POCT setting, detection was performed using an LFA. We evaluated the detection range of STACS by testing inactivated SARS-CoV-2 reference material across a wide RNA concentration range. As shown in Fig. 4C & 4D, STACS produced significantly elevated fluorescence signals for RNA inputs from 3.6 to 1006.1 copies per reaction compared with the negative control, indicating robust discrimination across more than two orders of magnitude. The consistent increase in signal with rising target concentration further supports the use of STACS for semi-quantitative analysis over this range.

Together, these results establish that STACS, when combined with a multiple-gRNA recognition strategy, enables rapid, amplification-free, and ultrasensitive detection of SARS-CoV-2 RNA with a broad dynamic range.

#### Effect of target mismatches on LbuCas13a and STACS activation

To evaluate mismatch tolerance and potential off-target signal generation, we first examined th response of LbuCas13a to N-gene RNA targets containing mismatches at different positions. Single-nucleotide mismatches produced strongly position-dependent effects on Cas13a activation. Relative to the fully matched N-gene target (1.00), M4, M8, M16, and M20 retained 0.80, 0.84, 0.68, and 0.70 of the fluorescence response, respectively. Introducing two mismatches resulted in substantially greater suppression, with M8+11 and M4+16 retaining only 0.34 and 0.08 of the fully matched response, respectively (Fig. S13). We next examined whether this mismatch-dependent discrimination was retained following downstream signal amplification in STACS. Using M16 and M4+16 as representative single- and double-mismatched targets, respectively, the background-corrected STACS signal decreased to 0.79 and 0.24 relative to the fully matched N-gene target (1.00), with both reductions being statistically significant (Fig. S14). These results demonstrate that mismatch-dependent differences in target recognition are retained in STACS rather than being completely masked by downstream Cas12a activation.

### 5. The detection performance of STACS for SARS-CoV-2 in environmental samples

The applicability of STACS for SARS-CoV-2 RNA detection in complex environmental matrices was evaluated using mud samples as a representative challenge. The workflow for environmental sample processing is illustrated in Fig. 5A. Briefly, mud samples were filtered to remove potentially inhibitory particulates, and subsequently the synthetic SARS-CoV-2 genomic RNA was tested by STACS. Before testing STACS, the activation ability of standard LbCas12a and LbuCas13a in 5% environmental mud sample was demonstrated (Fig. S8).

**Figure 5.**
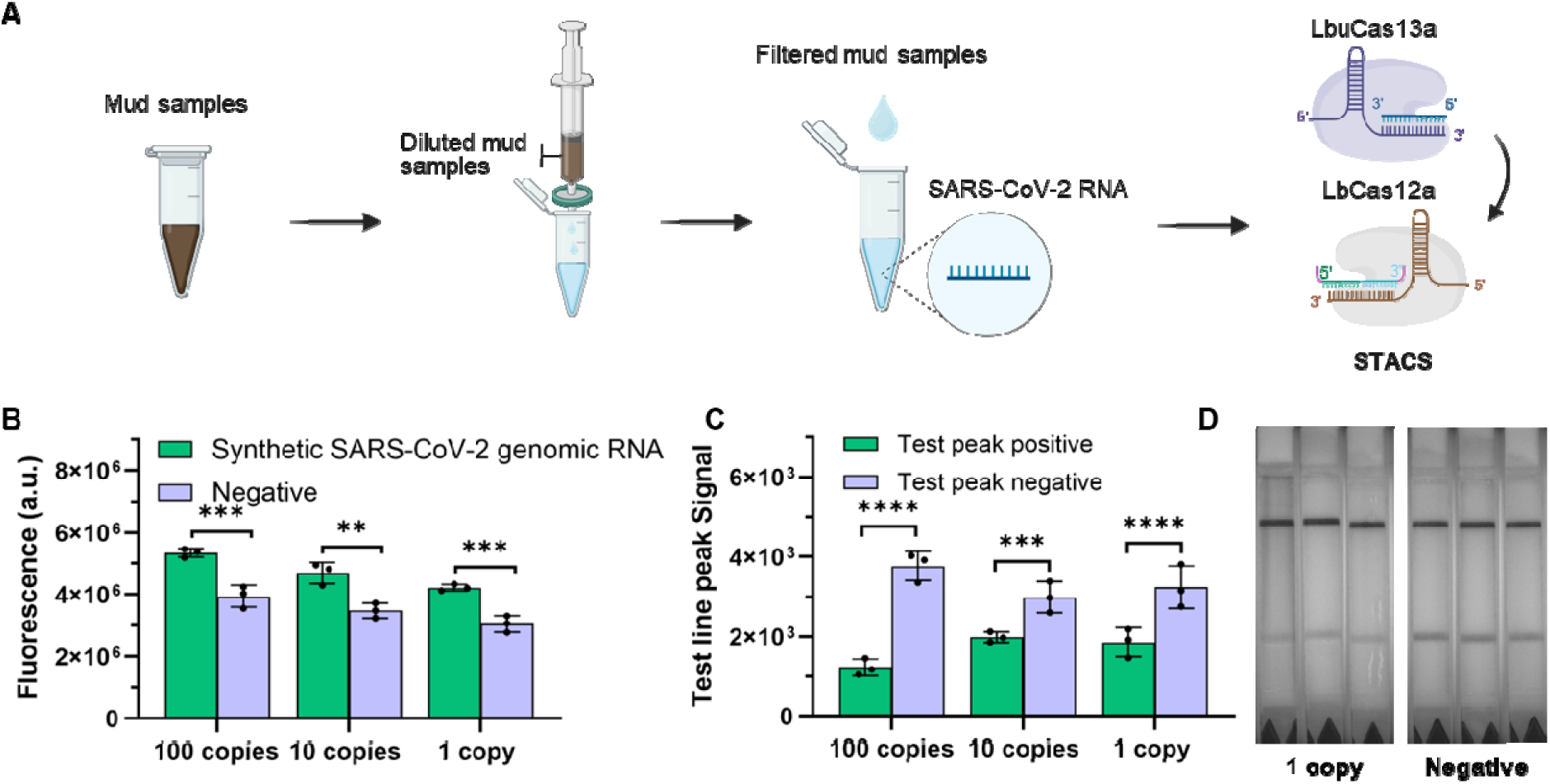
STACS application in environmental samples via fluorescent-based readout and LFA. (A). Schematic diagram of environmental mud sample preparation in STACS; (B) The detection ability of STACS for synthetic SARS-CoV-2 genomic RNA in 5% environmental mud sample via fluorescent-based readout; (C) The detection ability of STACS for synthetic SARS-CoV-2 genomic RNA in 5% environmental mud sample via LFA-based readout; (D) The corresponding LFA image of 1 copy in Figure C. n=3, error bars represent mean ± SD, * P ≤ 0.05, ** P ≤ 0.01, *** P ≤ 0.001, **** P ≤ 0.0001, ns = non-significant, a.u = arbitrary units.

Using fluorescence-based readout, STACS enabled clear discrimination between positive and negative samples in mud matrices. As shown in Fig. 5B, reactions containing 100, 10, and 1 copies of synthetic SARS-CoV-2 genomic RNA produced significantly higher fluorescence signals compared with negative controls, demonstrating that STACS retains high analytical sensitivity in environmental samples despite matrix complexity.

Consistent results were obtained using LFA readout. Quantitative analysis of test line peak intensities revealed statistically significant differences between positive and negative sample across all tested concentrations (Fig. 5C). Representative LFA strips further confirmed that STACS could reliably detect as few as 1 copy of synthetic SARS-CoV-2 genomic RNA in filtered mud samples (Fig. 5D & Fig. S9).

Collectively, these results demonstrate that STACS maintains robust performance in environmentally relevant matrices following minimal sample preparation. The ability to sensitively detect low-copy viral RNA in mud highlights the potential of STACS for environmental surveillance applications, such as wastewater monitoring and field-based pathogen detection.

### 6. Detection performance of STACS for SARS-CoV-2 in biological matrices

The performance of STACS in biological matrices was next evaluated using human serum as a representative sample. As illustrated in Fig. 6A, serum samples were pretreated with proteinase K followed by heat inactivation to mitigate matrix-associated interference prior to STACS analysis. The pretreatment effect of proteinase K was demonstrated to be highly effective (Fig. S10).

**Figure 6.**
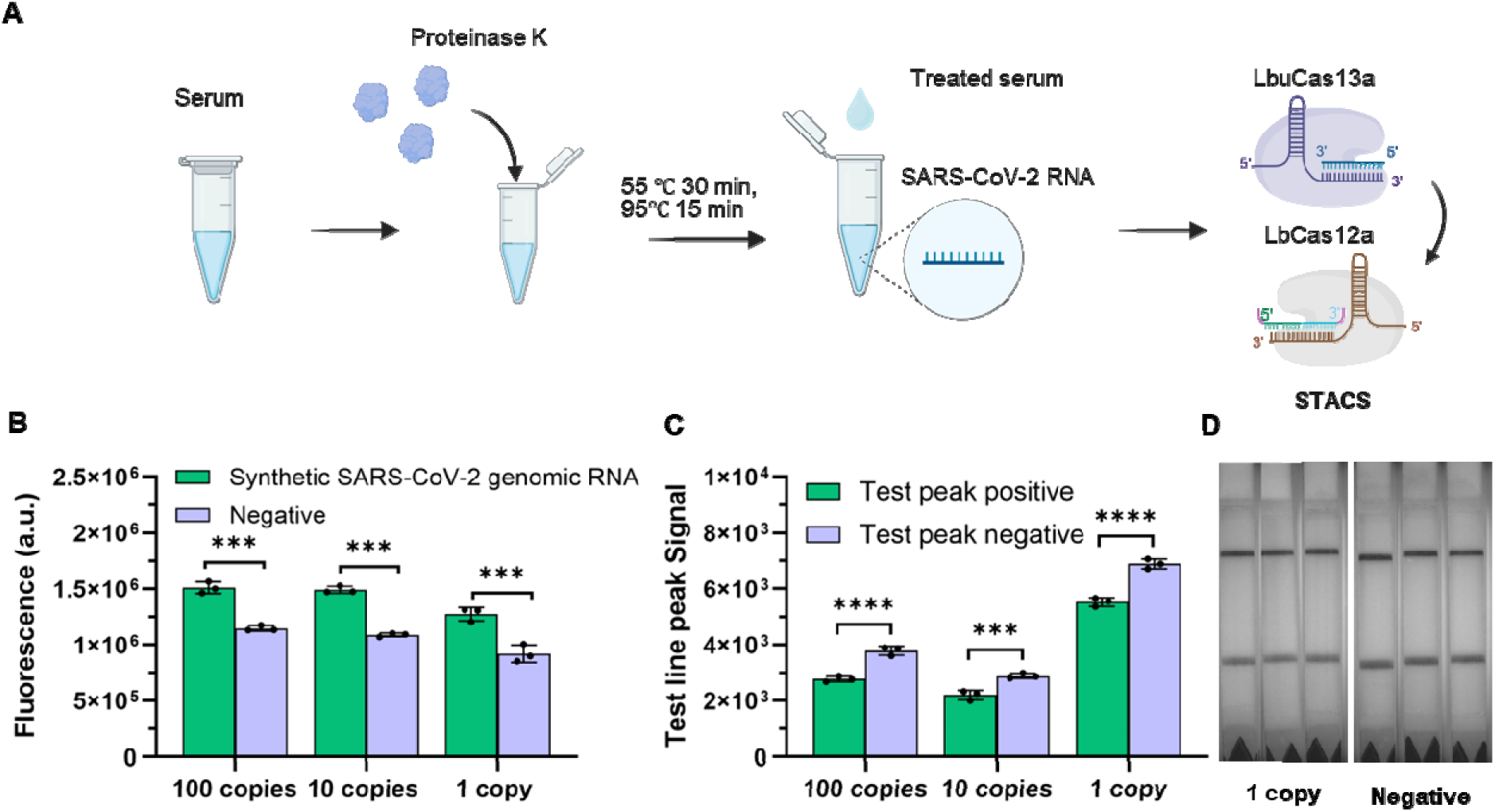
STACS application in 10% human serum via fluorescent-based readout and LFA. (A) Schematic diagram of human serum preparation in STACS; (B) The detection ability of STACS for 1-100 copies of synthetic SARS-CoV-2 genomic RNA in 10% human serum via fluorescent-based readout; (C) The detection ability of STACS for 1-100 copies of synthetic SARS-CoV-2 genomic RNA in 10% human serum via LFA readout (C); D. The corresponding images of 1 copy in Figure C. n=3, error bars represent mean ± SD, * P ≤ 0.05, ** P ≤ 0.01, *** P ≤ 0.001, ns = non-significant.

Using fluorescence-based readout, STACS enabled reliable detection of synthetic SARS-CoV-2 genomic RNA in 10% human serum. As shown in Fig. 6B, samples containing 100, 10, and 1 copies of synthetic SARS-CoV-2 genomic RNA produced significantly higher fluorescence signals than negative controls, demonstrating that STACS retains high sensitivity in serum despite the presence of endogenous biomolecules.

Consistent results were obtained using LFA readout. Quantitative analysis of test line peak intensities revealed clear and statistically significant discrimination between positive and negative samples across all tested concentrations (Fig. 6C). Representative LFA strips further confirmed that STACS could detect as few as 1 copy of synthetic SARS-CoV-2 genomic RNA in serum samples following proteinase K pretreatment (Fig. 6D & Fig. S11). These results demonstrate that STACS maintains robust amplification-free detection performance in clinically relevant serum matrices after simple enzymatic pretreatment, supporting its potential utility for rapid molecular diagnostics in real biological samples.

Encouraged by the robust performance of STACS in serum and environmental matrices, we next evaluated its applicability in human saliva, a clinically relevant and non-invasive sample type (Fig. 7A). To better simulate the processing of virus-containing biological samples, certified inactivated SARS-CoV-2 reference material was spiked into undiluted human saliva, followed by proteinase K treatment and heat treatment prior to STACS analysis. The activities of LbCas12a and LbuCas13a were first assessed in proteinase K-pretreated 10% human saliva, with no significant reduction observed compared with the corresponding controls (Fig. S12).

**Figure 7.**
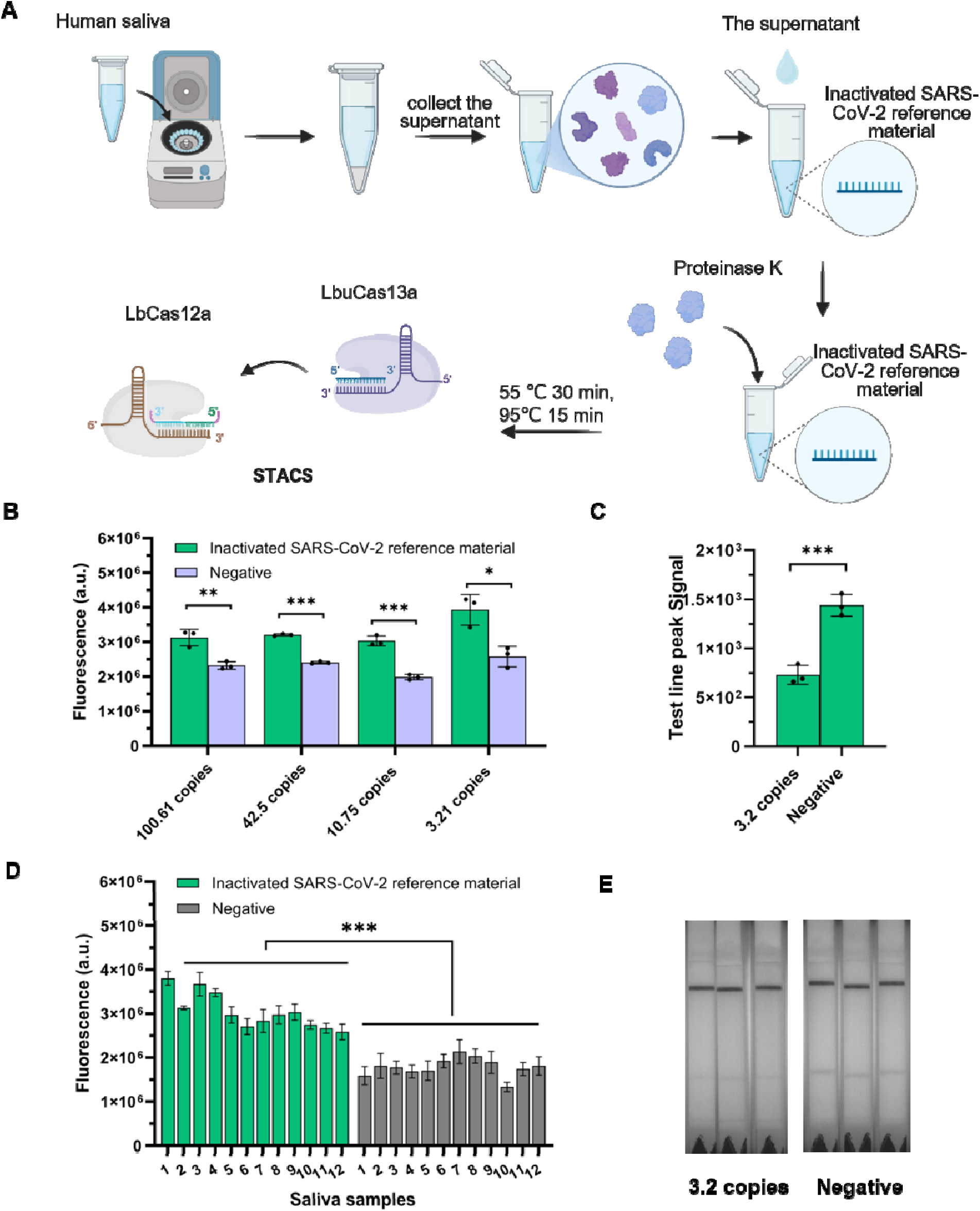
Detection of inactivated SARS-CoV-2 reference material in human saliva using STACS with fluorescence and LFA readouts. (A) Schematic of SARS-CoV-2-spiked human saliva sample processing and STACS detection. (B) Fluorescence-based detection of different concentrations of inactivated SARS-CoV-2 reference material following proteinase K treatment in human saliva. (C). LFA detection of inactivated SARS-CoV-2 reference material in processed human saliva. (D). Fluorescence-based detection of inactivated SARS-CoV-2 reference material corresponding to 3.21 copies per reaction across saliva samples from 12 independent donors. (E). Representative LFA detection of inactivated SARS-CoV-2 reference material corresponding to 3.21 copies per reaction across saliva samples from 12 independent donors following proteinase K treatment. n=3, error bars represent mean ± SD, * P ≤ 0.05, ** P ≤ 0.01, *** P ≤ 0.001, ns = non-significant.

Using fluorescence-based readout, STACS enabled sensitive detection of the inactivated SARS-CoV-2 reference material following processing in human saliva. Samples corresponding to 100.61, 42.5, 10.75, and 3.21 SARS-CoV-2 copies per reaction generated significantly higher fluorescence signals than negative controls (Fig. 7B), demonstrating amplification-free detection at low viral genome levels in the saliva matrix.

We further assessed the robustness of STACS across multiple individual saliva samples. Inactivated SARS-CoV-2 reference material corresponding to 3.21 copies per reaction was spiked into 12 independent human saliva samples and subjected to the same proteinase K and heat treatment. STACS consistently produced significantly higher fluorescence signals compared with corresponding negative controls (Fig. 7D), indicating good robustness against inter-individual sample variability in the saliva matrix.

Finally, we evaluated compatibility with a point-of-care LFA readout. Quantitative analysis revealed a significant difference in test-line peak signals between positive and negative saliva samples containing 3.21 SARS-CoV-2 copies per reaction (Fig. 7C), consistent with the representative LFA results obtained across individual saliva samples (Fig. 7E). Collectively, these results demonstrate that STACS enables sensitive, amplification-free detection of SARS-CoV-2 reference material in human saliva following minimal sample processing, supporting its potential for further development toward non-invasive and point-of-care molecular diagnostics.

## Discussion

In this study, we establish the molecular principles that define how fragmented nucleic acid triggers activate Cas12a and show how these principles collectively enable an architecture that expands a single target recognition event into large-scale downstream Cas RNP activation. Our mechanistic analyses reveal three fundamental requirements for Cas12a activation by split triggers: (i) external nucleotide extensions are tolerated, whereas internal insertions disrupt spacer complementarity and abolish R-loop formation; (ii) activation strictly requires two physically independent fragments, as covalent linkage prevents the conformational flexibility needed for stepwise R-loop propagation; and (iii) efficient activation depends on precise fragment lengths, with a short seed-nucleating strand and a complementary R-loop extending strand providing the optimal synergistic configuration. These insights resolve the previously undefined sequence and structural rules governing split-trigger compatibility with Cas12a.

By mapping these mechanistic constraints, we demonstrate why LbuCas13a cleavage naturally produces mechanistically optimal activators - short, independent DNA fragments bearing external overhangs - which directly feed into Cas12a activation. Notably, the efficiency of LbuCas13a-mediated cleavage of the linear mediator represents a critical determinant of STACS performance, as it governs the rate and abundance of split-trigger generation required for downstream Cas12a activation. This coincidence enables a predictable, sequential signal relay in which Cas13-based RNA recognition is immediately converted into Cas12a-mediated DNA signal amplification without any enzymatic preamplification. The optimized split-trigger pair (11 nt + 10 nt) maximizes the R-loop seeding efficiency and underpins the high analytical sensitivity of the system.

Leveraging these design principles, we developed STACS, a linear mediator-powered Cas13-Cas12 tandem platform that achieves amplification-free detection down to 1 copy/µL within 15 minutes. Unlike conventional CRISPR assays(19–21), which rely on single-enzyme *trans*-cleavage, STACS decouples recognition and reporting into two orthogonal nucleases. This design enhances signal gain, suppresses background activation, and ensures that Cas12a activity occurs only when triggered by bona fide Cas13 cleavage products. As a result, STACS enables highly sensitive nucleic acid detection without thermal cycling or isothermal amplification, substantially simplifying workflow and eliminating the risk of contamination.

Importantly, the robustness of the STACS linear mediators translates into strong performance across diverse sample matrices, including human serum, human saliva, and environmental mud. Proteinase K treatment effectively stabilizes split mediators in biological fluids, while simple filtration restores CRISPR activity in environmental samples. Across all matrices, STACS consistently detected SARS-CoV-2 genomic RNA at low copy numbers, demonstrating its utility for point-of-care diagnostics, field surveillance, and rapid testing in resource-limited settings. Although this study employs certified SARS-CoV-2 reference materials with traceable quantification and demonstrates robust performance across complex biological and environmental matrices, the current results primarily establish analytical validity. Further work is required to achieve large-scale clinical validation, direct benchmarking against gold-standard methods such as RT-qPCR, and systematic evaluation of stability and deployment readiness. Future studies will focus on validating STACS using authentic clinical specimens to confirm its clinical applicability. In addition, while STACS exhibits concentration-dependent signal output, its nonlinear amplification driven by sequential Cas activation positions it primarily as a highly sensitive qualitative detection platform with limited semi-quantitative capability.

More broadly, our mechanistic framework provides a foundation for rationally engineering muti-stage CRISPR systems (Table S1) (13,17,22–26). The rules identified here for Cas12a-including extension position, fragment independence, and length-dependent R-loop dynamics-provide design principles for split-trigger-mediated Cas12a activation and may also inform future investigation of analogous activation strategies in other CRISPR-Cas effectors, such as Cas13, Cas14, and other emerging effectors. Because split-trigger activation enables modular signal gating and combinatorial logic (12), STACS also offers opportunities for multiplexed sensing, conditional activation circuits, and high-fidelity molecular computation.

Overall, this work bridges a critical mechanistic gap in Cas activation biology and demonstrates how fundamental insights into R-loop initiation and propagation can yield practical, high-performance diagnostic technologies. The linear mediator-powered Cas13-Cas12 system represents a versatile platform for rapid, ultralow-input, amplification-free RNA/DNA detection suitable for clinical diagnostics, non-invasive liquid biopsy, environmental pathogen monitoring, and next-generation point-of-care testing.

## Conclusion

In this work, we define the molecular rules that govern Cas12a activation by fragmented nucleic acids and demonstrate how these principles can be exploited to engineer a programmable Cas13-Cas12 tandem detection system. We reveal that Cas12a activation depends critically on extension position, fragment independence, and optimized trigger length - mechanistic constraints that together determine whether split activators can successfully nucleate and propagate the R-loop. These insights explain why Cas13-mediated cleavage naturally produces functional Cas12a activators and provide a rational framework for designing future muti-stage CRISPR sensing systems.

Building on these mechanistic foundations, we developed STACS, an amplification-free Cas13-Cas12 system capable of detecting RNA targets at 1 copy/µL within 15 minutes. The platform maintains high analytical sensitivity across clinically and environmentally relevant matrices, including serum, saliva, and mud, and is compatible with both fluorescence and lateral-flow readouts. By decoupling RNA recognition from DNA reporting through a linear DNA-RNA-DNA mediator, STACS integrates mechanistic precision with practical robustness, enabling ultrafast and field-deployable molecular diagnostics. This study bridges a key mechanistic gap in CRISPR activation biology and introduces a versatile architecture based on sequential Cas activation that supports rapid, low-input RNA/DNA detection for point-of-care testing, environmental surveillance, and translational biomedical applications.

## Acknowledgements

The authors acknowledge the support of the Australia’s Economic Accelerator (AEA) (AE240300106) of FD. and the ARC Discovery funding (DP240103024) to EMG and FD.

## Author contributions

Rui Sang: conceptualisation, investigation, methodology, formal analysis, writing – original draft, writing – review & editing. Avin Mohammadnejad Mavaneh: investigation, methodology, writing – review & editing. Ewa M. Goldys: writing – review & editing, supervision, funding acquisition, project administration. Fei Deng: conceptualisation, methodology, formal analysis, writing – original draft, writing – review & editing, supervision, funding acquisition, project administration.

