## Supplemental file for "Modular split-trigger synergistic activation enables a programmable Cas13-12 system"

### Supplementary Methods:

#### Investigation of trigger mechanism of split mediator for Lbcas12a RNP via LFA

To investigate split trigger activation mechanism for LbCas12a via LFA, 2μL of 1μM of trigger (Trigger 1, Trigger 3, Trigger 1 & Trigger 2 and Intact Trigger) was added into 98 μL of standard CRISPR/ LbCas12a reaction solution (20nM of LbCas12a protein, 20nM of gRNA, 166nM of 8C reporter, and 1mL of rCutSmart buffer). The reaction mixture was incubated at 37 °C for 30 min. Then, 6 μL of reaction solution were mixed with 94 μL of Milenia running buffer at 30 min, followed by adding Milenia HybriDetect 1 strip into the tube and allowed to run for around 3 minutes. Lastly, strips were read on a lateral Flow Reader (AX-2X-S, Axxin Pty Ltd, Australia).

#### Assessment of LbuCas13a *trans*-cleavage toward the linear mediator using different length of linker

LbuCas13a ribonucleoprotein complexes were prepared by mixing 2.5 μL of 20 μM LbuCas13a endonuclease with 2.5 μL of 20 μM LbuCas13a SARS-CoV-2 gRNA in 1.25 mL of 1× rCutSmart buffer. The resulting RNP mixture was evenly divided into five aliquots (250 μL per tube). To each aliquot, 2.5 μL of 100 μM linear mediator reporters with different linker lengths (split 1-2, split 1-2-1U, split 1-2-3U, split 1-2-5U and split 1-2-7U) was added. The prepared Cas13a reaction mixtures were stored at 4 °C prior to use, with 50 μL reserved as a no-target negative control. To initiate Cas13a activation, 1 μL of 10 μM SARS-CoV-2 RNA was added to 200 μL of each prepared reaction mixture. Reactions were incubated at 37 °C, and 50 μL aliquots were collected at 15, 30, 60, and 90 min. Enzymatic activity was terminated by heating the collected samples at 95 °C for 10 min. These collected reaction solutions were checked by agarose gel electrophoresis. Briefly, 5% of the agarose gels were prepared in 1× TAE buffer and stained with SYBR Gold DNA dye. 10 μL of samples were premixed with 2 μL of 6X DNA gel loading dye and then loaded into gel for electrophoresis, which was carried out for 60 min at a constant voltage of 100V. 3 μL of 10 bp DNA ladder was used for molecular weight reference. Gel images were visualized by using Gel Doc + XR image system (Bio-Rad Laboratories Inc., USA).

#### Assessment of LbuCas13a trans-cleavage on the linear mediator (split 2-1-10-11-5U-F) using fluorescence readout

LbuCas13a: 1 μL of 20 μM of LbuCas13a endonuclease and 1 μL of 20 μM of LbuCas13a SARS-CoV-2 gRNA were mixed at 1 mL of 1× rCutSmart, followed by addition of 3 μL of 10 μM of split 2-1-10-11-5U-F reporter. The prepared CRISPR/Cas13a reaction mixture was preincubated for 30 min at 37°C and then 10 μL of 200 nM or 10 nM of SARS-CoV-2 N gene RNA or DPBS (as negative control) was added into 90 μL of CRISPR/LbuCas13a reaction solution. The reaction was carried out at 37 °C, and the fluorescence intensity at Ex/Em of 570/615 nm was determined by using a plate reader (iD5 Spectramax, Molecular Devices, USA).

**Optimization of reaction buffers**

**LbCas12a:** 0.5 μL of 20 μM of LbCas12a endonuclease and 0.5 μL of 20 μM of intact trigger gRNA were mixed at 1 mL of different reaction buffers (1× rCutSmart/ NEB r2.1/ NEB 3.1/ NEB 4 buffer), followed by addition of 5 μL of 100 μM of 6C reporter. The prepared CRISPR/Cas12a reaction mixture was preincubated for 30 min at 37°C and then 2 μL of 1 μM of trigger (trigger 1, trigger 2, trigger 1&trigger 2, split 2-1 or split 1-2 s) was added into 98 μL of CRISPR/LbCas12a reaction solution. The reaction was carried out at 37 °C, and the fluorescence intensity at Ex/Em of 570/615 nm was determined by using a plate reader (iD5 Spectramax, Molecular Devices, USA).

**LbuCas13a:** 0.5 μL of 20 μM of LbuCas13a endonuclease and 0.5 μL of 20 μM of LbuCas13a SARS-CoV-2 gRNA were mixed at 1 mL of the indicated buffers (1× rCutSmart/ NEB r2.1/ NEB 3.1/ NEB 4 buffer), followed by addition of 5 μL of 100 μM of 5U reporter. The prepared CRISPR/Cas13a reaction mixture was preincubated for 30 min at 37°C and then 2μL of 1μM of SARS-CoV-2 N gene RNA or DPBS (as negative control) was added into 98 μL of CRISPR/LbuCas13a reaction solution. The reaction was carried out at 37 °C, and the fluorescence intensity at Ex/Em of 570/615 nm was determined by using a plate reader (iD5 Spectramax, Molecular Devices, USA).

**Activation efficiency assessment of different gRNAs for LbuCas13a using the corresponding SARS-CoV-2 RNA targets**

To evaluate the activation efficiency of different gRNAs for LbuCas13a, 4 μL of 20 μM LbuCas13a endonuclease and 4 μL of 20 μM LbuCas13a SARS-CoV-2 gRNA targeting the N, M, E or ORF genes was mixed at 1 mL 1× rCutSmart, followed by addition of 1.66 μL of 100 μM 5U reporter. For each CRISPR/Cas13a *trans*-cleavage activation reaction, 2 μL of 1 μM SARS-CoV-2 RNA corresponding to the N, M, E, or ORF gene was added into 98 μL prepared reaction buffer. The fluorescence intensity at Ex/Em of 570/615 nm was determined by using a plate reader (iD5 Spectramax, Molecular Devices, USA) at 37 °C.

**Effect of step-one reaction time on STACS fluorescence signal**

To evaluate the influence of step-one reaction time on the STACS fluorescence response to reference SARS-CoV-2 RNA standards, 10 μL of 360 copies/mL of reference SARS-CoV-2 RNA was added into 90 μL of standard multiple gRNA CRISPR/LbuCas13a reaction solution. The reaction mixture was incubated at 37 °C for 15 min, 5 min and 30 second, followed by adding 10 μL of 10 times standard multiple gRNA CRISPR/LbCas12a reaction solution. The fluorescence intensity at Ex/Em of 570/615 nm was determined by using a plate reader (iD5 Spectramax, Molecular Devices, USA) at 37 °C.

**Stability study of STACS in environmental samples**

To assess whether environmental mud samples induce nonspecific activation of CRISPR/LbuCas13a or CRISPR/LbCas12a systems, standard reaction solutions were prepared. The CRISPR/LbuCas13a reaction solution contained 40 nM of LbuCas13a protein, 40 nM of gRNA, and 500 nM of 5U reporter in 1 mL of 1× rCutSmart buffer. The CRISPR/LbCas12a reaction solution contained 40 nM of LbCas12a protein, 40 nM of gRNA, and 500 nM of 6C reporter in 1 mL of 1× rCutSmart buffer. For target-free assessment, 10 μL of 5% (v/v) environmental mud was added to 90 μL of the prepared reaction solutions. Fluorescence signals were recorded at an excitation/emission wavelength of 570/615 nm using a microplate reader (SpectraMax iD5, Molecular Devices, USA) at 37 °C.

To evaluate the effect of environmental mud on target-induced Cas activation, 10 μL of 5% environmental mud or filtered 5% environmental mud was mixed with 90 μL of the standard CRISPR/LbuCas13a or CRISPR/LbCas12a reaction solutions (40 nM of RNP and 500 nM of reporter in 1× rCutSmart buffer). Cas activation was initiated by adding 10 μL of 1 nM of SARS-CoV-2 RNA or intact trigger, respectively. Fluorescence measurements were performed at 37 °C using a microplate reader (SpectraMax iD5, Molecular Devices, USA) with an excitation/emission wavelength of 570/615 nm.

**The stability study of Split 2-1-11-10-5U-F in human serum**

To investigate the stability of Split 2-1-11-10-5U-F in human serum, 60 nM of Split 2-1-11-10-5U-F were mixed with 1x rCutSmart buffer containing 10% human serum without or with Proteinase K pretreatment. The fluorescence intensity at Ex/Em of 570/615 nm was determined by using a plate reader (iD5 Spectramax, Molecular Devices, USA) at 37 °C.

**The stability study of STACS in human saliva**

To assess whether human saliva induces nonspecific activation of CRISPR/LbuCas13a and CRISPR/LbCas12a systems, 10 μL of 10% (v/v) human saliva with or without Proteinase K pretreatment, was added to 90 μL of standard CRISPR/LbuCas13a reaction solution (40 nM of LbuCas13a protein, 40 nM of gRNA, and 500 nM of 5U reporter in 1× rCutSmart buffer) or standard CRISPR/LbCas12a reaction solution (40 nM of LbCas12a protein, 40 nM of gRNA, and 500 nM of 6C reporter in 1× rCutSmart buffer). The fluorescence intensity at Ex/Em of 570/615 nm was determined by using a plate reader (iD5 Spectramax, Molecular Devices, USA) at 37 °C.

To investigate the Proteinase K pretreated human saliva samples effects for CRISPR/LbuCas13a and CRISPR/LbCas12a reaction after targets activation, 10 μL of 10 nM of reference SARS-CoV-2 Standards RNA or Intact Trigger in 10% proteinase k pretreated human saliva were mixed with 90 μL of standard CRISPR/LbuCas13a or CRISPR/LbCas12a solutions (40 nM of RNP and 500nM of reporter in 1mL of rCutSmart buffer). The fluorescence intensity at Ex/Em of 570/615 nm was determined by using a plate reader (iD5 Spectramax, Molecular Devices, USA) at 37 °C.

**Mismatch sensitivity of LbuCas13a toward single- and double-mismatched RNA targets**

LbuCas13a ribonucleoprotein (RNP) complexes were prepared by mixing 1 µL of 20 µM LbuCas13a endonuclease with 1 µL of 20 µM LbuCas13a SARS-CoV-2 N-gene gRNA in 1 mL of 1× rCutSmart buffer, followed by the addition of 1.66 µL of 100 µM 5U reporter. The LbuCas13a reaction mixture was preincubated at 37 °C for 30 min and stored at 4 °C prior to use. For mismatch sensitivity assessment, 10 µL of 10 nM synthetic SARS-CoV-2 N-gene RNA containing either the fully matched sequence or the indicated single or double nucleotide substitutions was added to 90 µL of the LbuCas13a reaction mixture. DPBS was used as the negative control. Reactions were incubated at 37 °C, and fluorescence was monitored at excitation/emission wavelengths of 570/615 nm using a SpectraMax iD5 microplate reader (Molecular Devices, USA).

**Mismatch sensitivity assessment of STACS**

The mismatch sensitivity of STACS was evaluated using the fully matched SARS-CoV-2 N-gene RNA and mismatched RNA targets M16 and M4+16, containing a single nucleotide substitution and two nucleotide substitutions, respectively. Briefly, 10 µL of 100 aM synthetic N-gene, M16, or M4+16 RNA was added to 90 µL of the standard multiple-gRNA LbuCas13a reaction mixture to initiate LbuCas13a trans-cleavage. After incubation at 37 °C for 30 min, 10 µL of 10× standard LbCas12a reaction mixture was added to initiate downstream Cas12a activation. The reactions were maintained at 37 °C, and fluorescence was monitored at excitation/emission wavelengths of 570/615 nm using a SpectraMax iD5 microplate reader (Molecular Devices, USA).

**Supplementary Figures:**

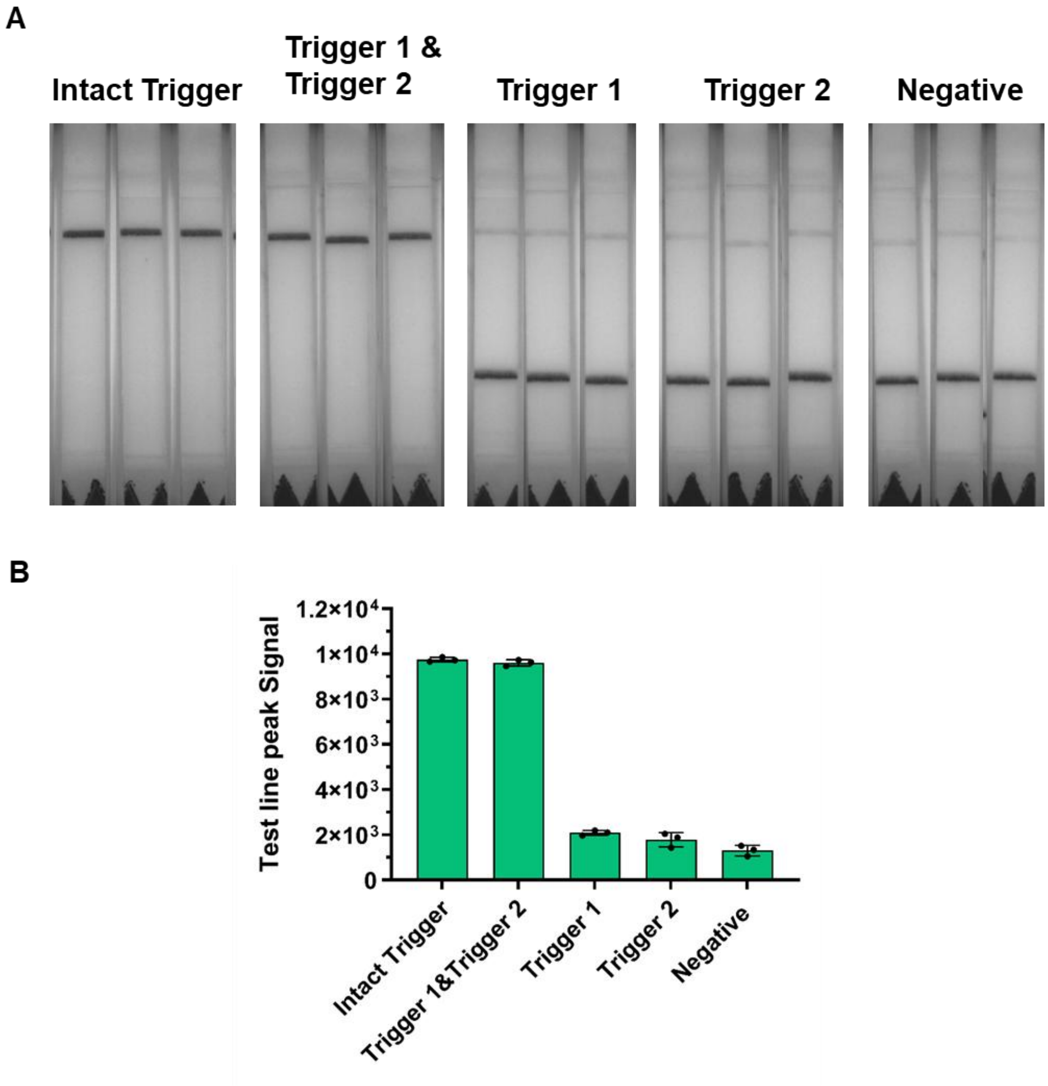


**Figure S1**. (A) Split triggers are able to fully activate LbCas12a using LFA; (B) the corresponding upper line peak analysis of figure A.

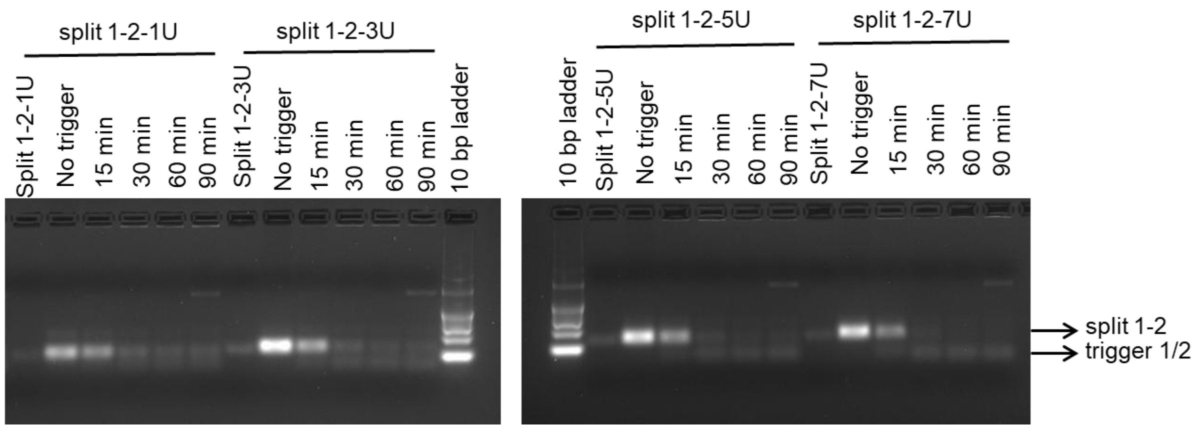


**Figure S2**. Assessment of LbuCas13a *trans*-cleavage on the linear mediator using agarose gel electrophoresis analysis (100V 30 min). The linear mediator was successfully cleaved by activated LbuCas13a.

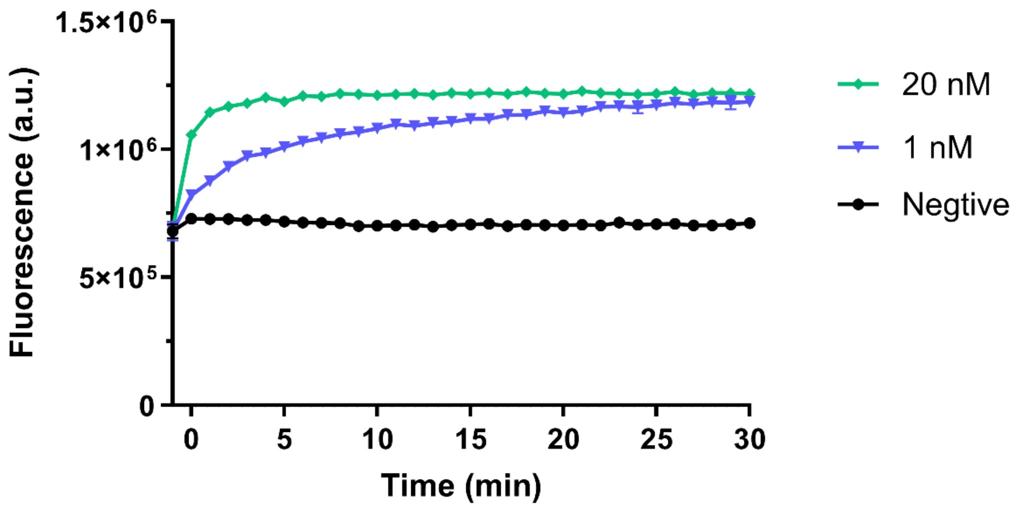


**Figure S3**. Assessment of LbuCas13a trans-cleavage on the linear mediator (split 2-1-10-11-5U-F) using fluorescence readout (N gene RNA as target).

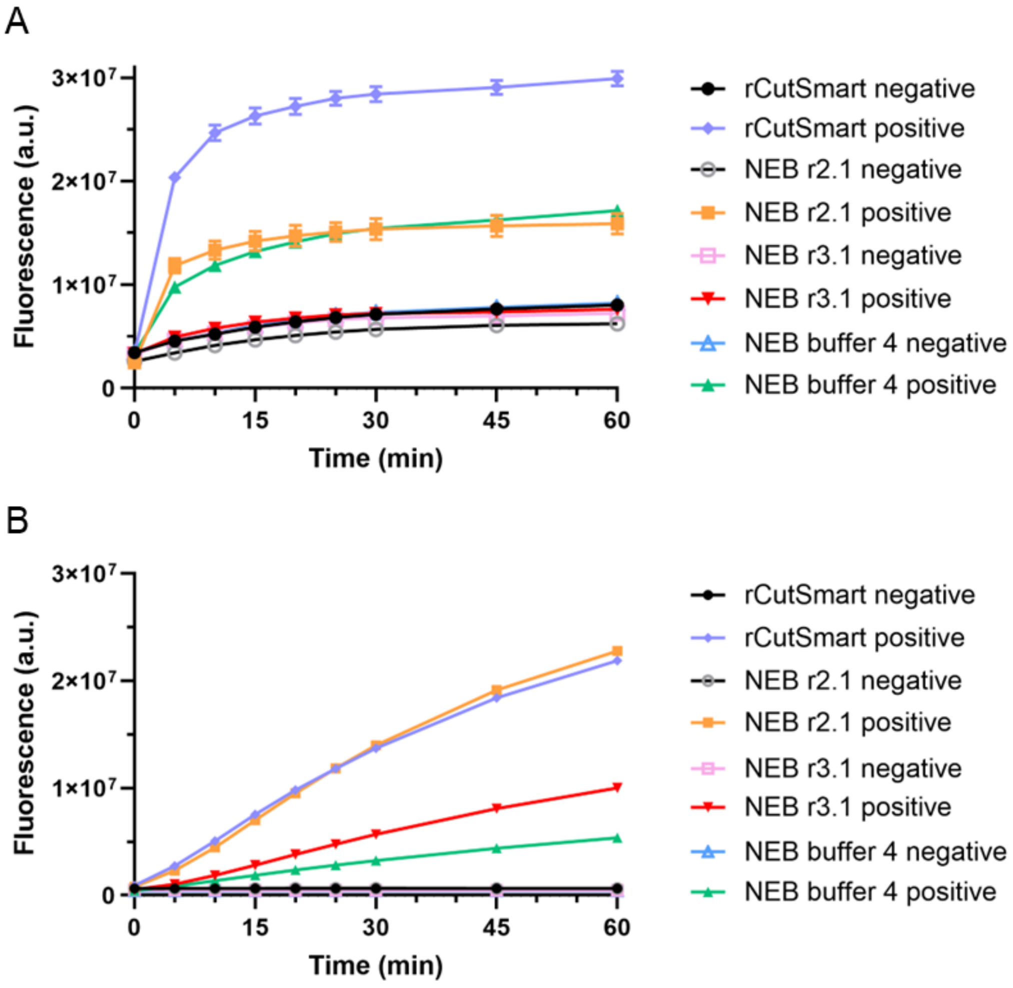


**Figure S4**. The reaction buffers optimization for LbuCas13a activated by SARS-CoV-2 RNA (A) and LbCas12a activated by Intact trigger (B) at 37°C. The optimum buffer for STACS was found to be rCutSmart buffer. n=3, error bars represent mean ± SD, * $P \leq 0.05$, ** $P \leq 0.01$, *** $P \leq 0.001$, **** $P \leq 0.0001$, ns = non-significant, a.u = arbitrary units.

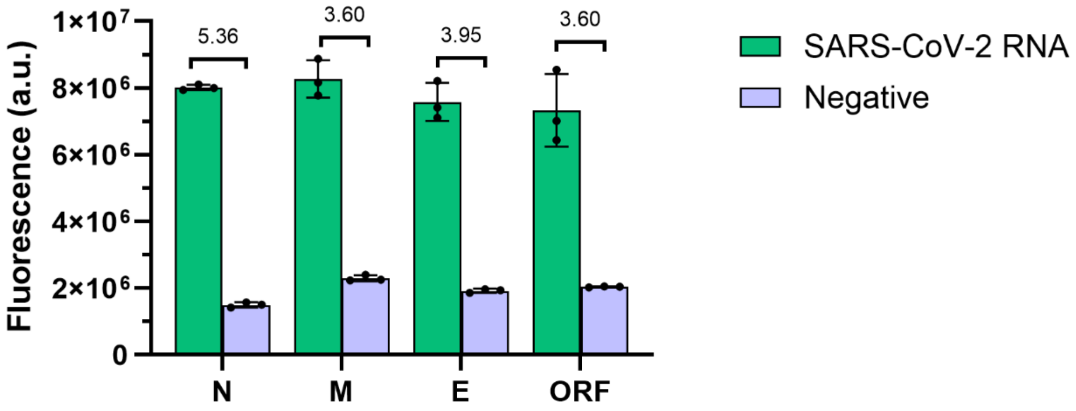


**Figure S5**. Assessment of the activation efficiency of different gRNAs for LbuCas13a using the corresponding SARS-CoV-2 RNA targets. n=3, error bars represent mean ± SD, * P ⩽ 0.05, ** P ⩽ 0.01, *** P ⩽ 0.001, **** P ⩽ 0.0001, ns = non-significant, a.u = arbitrary units.

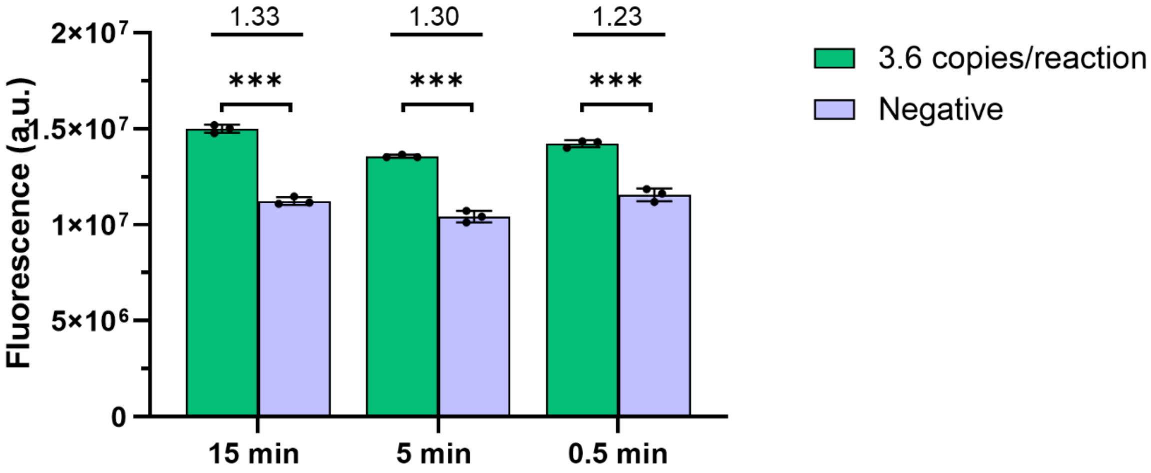


**Figure S6**. The fluorescence signal of STACS for 3.6 copies reference SARS-CoV-2 Standards RNA with different reaction time for step one. Since the incubation for step two was 15 min, thus, the total incubation time of STACS was optimized to be close to 15min. n=3, error bars represent mean ± SD, * P ⩽ 0.05, ** P ⩽ 0.01, *** P ⩽ 0.001, **** P ⩽ 0.0001, ns = non-significant, a.u = arbitrary units.

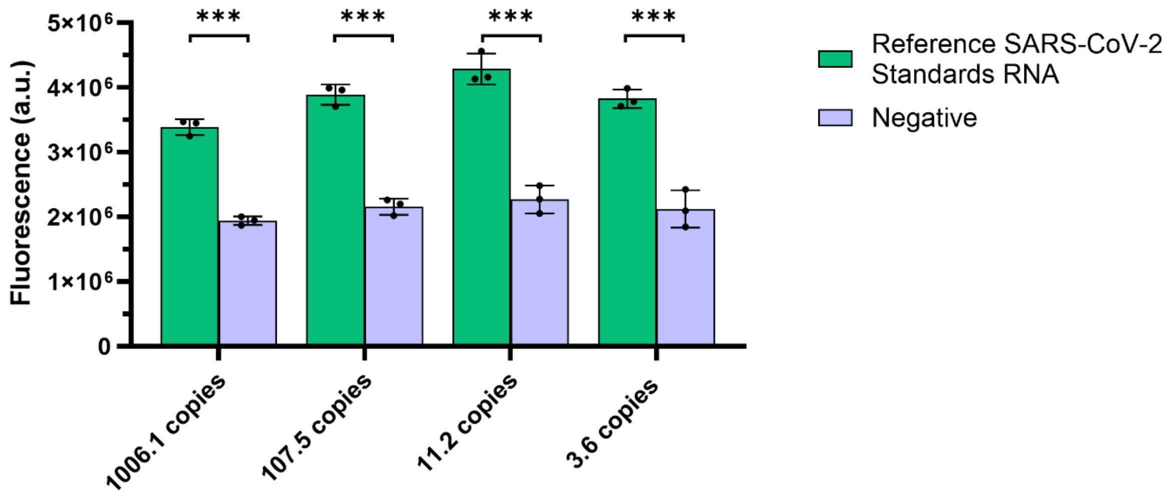


**Figure S7**. Fluorescence-based detection of reference SARS-CoV-2 RNA standards using the STACS platform with a multiple gRNA strategy targeting the N, M, E, and ORF genes. n=3, error bars represent mean ± SD, * $P \leqslant 0.05$, ** $P \leqslant 0.01$, *** $P \leqslant 0.001$, **** $P \leqslant 0.0001$, ns = non-significant, a.u = arbitrary units.

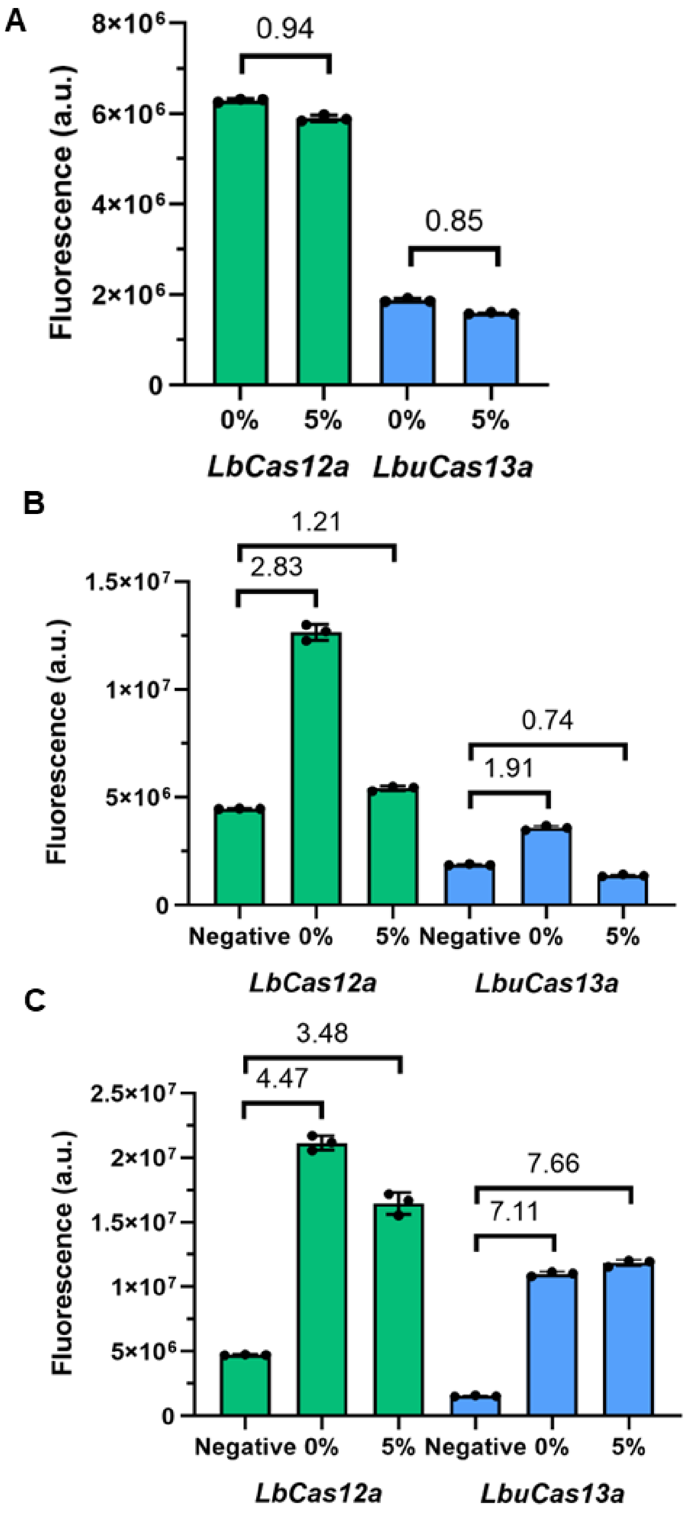


**Figure S8**. Influence of environmental mud on LbCas12a and LbuCas13a reaction performance. (A). Stability of LbCas12a/LbuCas13a RNPs and reporters in the presence of 5% environmental mud under target-free conditions. Environmental mud alone did not activate LbCas12a/LbuCas13a RNPs and induced instability of the 5U/6C reporters; (B). Effect of environmental mud on target-dependent activation of LbCas12a and LbuCas13a. Reactions containing 5% environmental mud were tested with 100 pM SARS-CoV-2 RNA or 1 nM

Intact trigger, respectively, showing significant suppression of Cas activation; (C). Effect of filtration on mud-induced interference. Filter-treated 5% environmental mud did not significantly affect LbCas12a or LbuCas13a reactions in the presence of 100 pM SARS-CoV-2 RNA or 1 nM Intact trigger; n=3, error bars represent mean ± SD, * $P \leqslant 0.05$, ** $P \leqslant 0.01$, *** $P \leqslant 0.001$, **** $P \leqslant 0.0001$, ns = non-significant, a.u = arbitrary units.

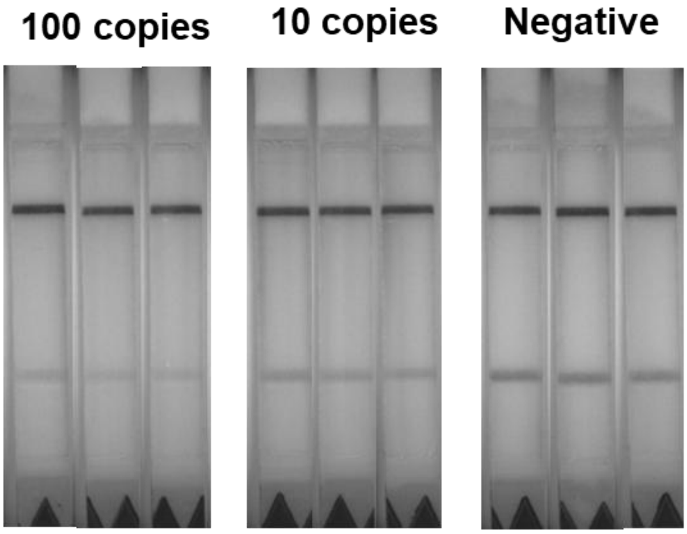


**Figure S9**. Detection of 100-10 copies of synthetic SARS-CoV-2 genomic RNA in 5% environmental mud using the STACS platform with a multiple gRNA strategy via LFA readout. Each condition was tested in triplicate (n = 3).

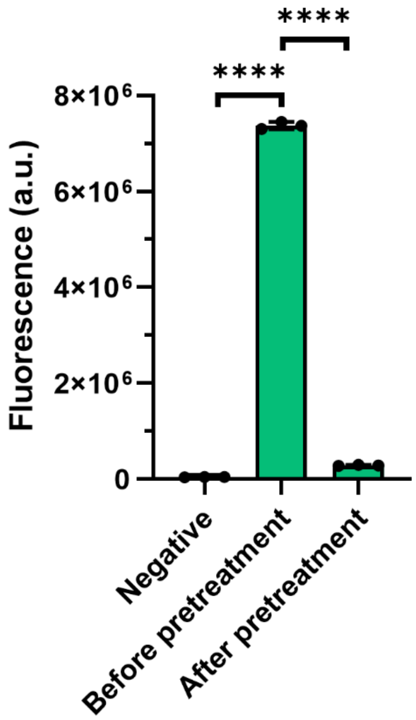


**Figure S10**. Stability assessment of split 2-1-11-10-5U-F (60 nM) in 10% human serum with and without Proteinase K pretreatment. The split 2-1-11-10-5U-F was very stable in Proteinase K treated human serum sample. n=3, error bars represent mean ± SD, * P ⩽ 0.05, ** P ⩽ 0.01, *** P ⩽ 0.001, **** P ⩽ 0.0001, ns = non-significant, a.u = arbitrary units.

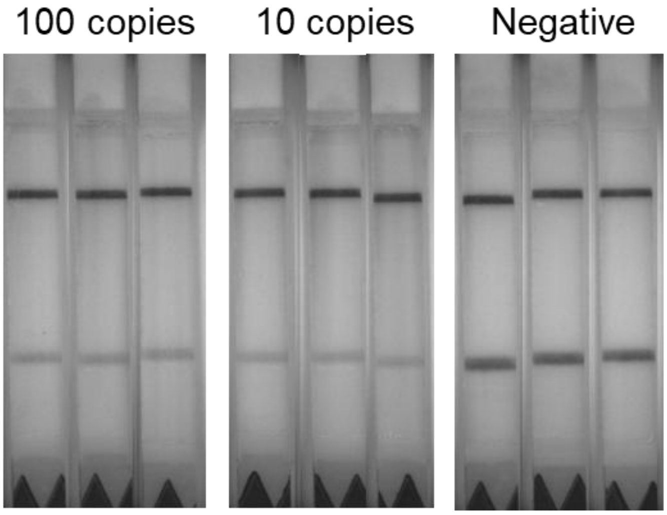


**Figure S11**. Detection of 100-10 copies of genomic SARS-CoV-2 RNA in 10% (v/v) human serum using the STACS platform with a multiple gRNA strategy via LFA readout. Each condition was tested in triplicate (n = 3).

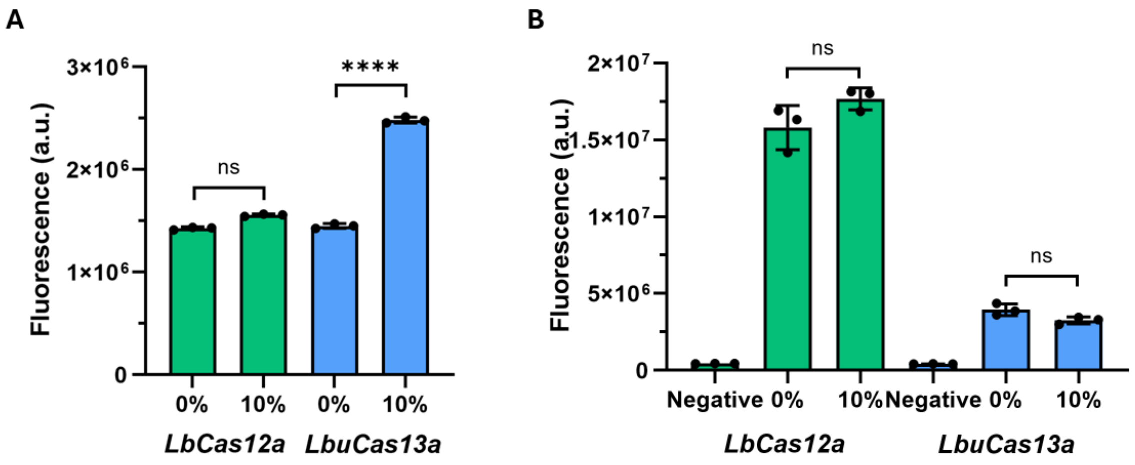


**Figure S12**. Influence of human saliva on LbCas12a and LbuCas13a reaction performance. (A). Stability of LbCas12a/LbuCas13a RNPs and reporters in the presence of 10% human saliva without Proteinase K pretreatment under target-free conditions. Human saliva alone did not induce activation of LbCas12a RNPs or instability of the 6C reporter but triggered instability of the 5U reporter of LbuCas13a. (B). Effect of Proteinase K pretreatment on saliva-induced interference. Proteinase K-treated 10% human saliva did not significantly affect LbCas12a and LbuCas13a reactions. n=3, error bars represent mean ± SD, * $P \leqslant 0.05$, ** $P \leqslant 0.01$, *** $P \leqslant 0.001$, **** $P \leqslant 0.0001$, ns = non-significant, a.u = arbitrary units.

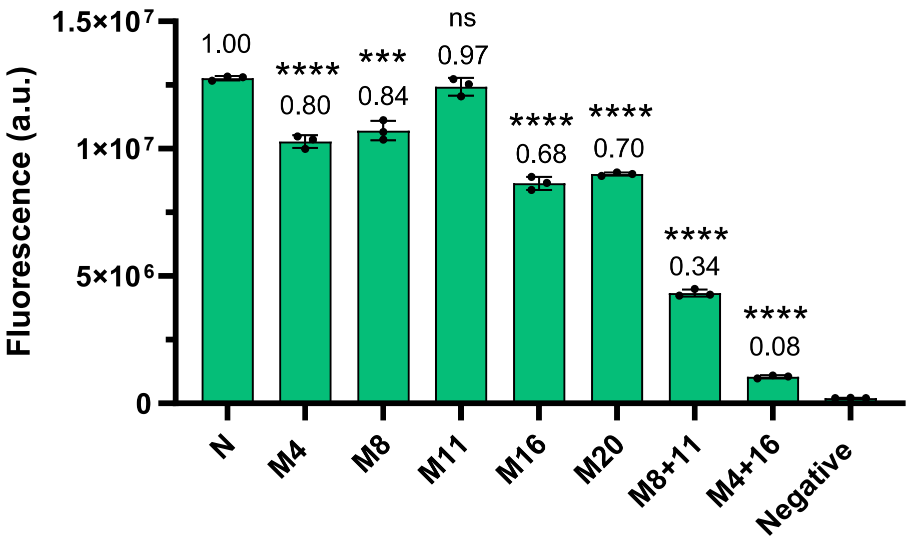


**Figure S13.** Mismatch sensitivity of LbuCas13a toward SARS-CoV-2 N-gene RNA targets. Fluorescence responses of LbuCas13a to the fully matched N-gene target (N), single-mismatched targets (M4, M8, M11, M16, and M20), double-mismatched targets (M8+11 and M4+16), and the negative control. Values above the bars indicate the relative fluorescence response normalized to that of the fully matched target (N = 1.00). n=3, error bars represent mean ± SD, *** $P \leq 0.001$, **** $P \leq 0.0001$, ns = non-significant, a.u = arbitrary units.

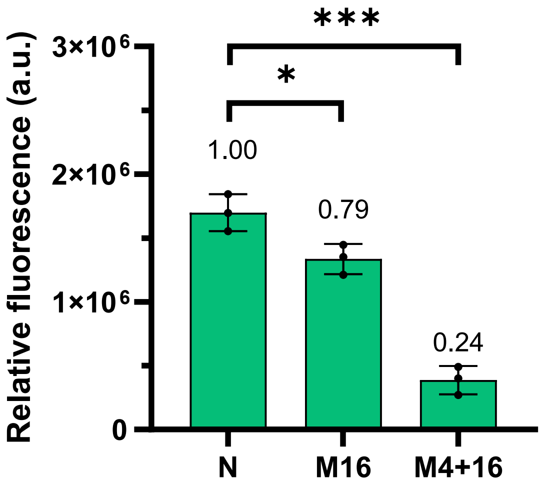


**Figure S14.** Effect of target mismatches on STACS activation. Fluorescence responses of STACS to the fully matched SARS-CoV-2 N-gene RNA target (N), the single-mismatched target M16, and the double-mismatched target M4+16. Fluorescence signals were background-corrected by subtracting the corresponding negative-control signal. Values above the bars indicate the relative fluorescence response normalized to that of the fully matched N-gene target (N = 1.00). n=3, error bars represent mean ± SD, * $P \leq 0.05$, *** $P \leq 0.001$, ns = non-significant, a.u = arbitrary units.

**Table S1**. The comparison with previously published tandem system.

| System | Enzyme Type | Mediator Structure/ Strategy | Detection Range / LOD | Sample Types | Detection Time | Reference |
|---|---|---|---|---|---|---|
| FIND-IT | Cas13a-Csm6 | Csm6 activator oligoadenylates $A_4$-$U_6$ | 31 copies/μL (~fM level) | SARS-CoV-2 in clinical respiratory swabs | ~20 min | (1) |
| casCRISPR | Cas13a-Cas14a | hairpin structured DNA oligonucleotides ST-HP with unpaired rU | ~1.33 fM | miRNA (miR-17) in Serum and cell extracts | 60 min | (2) |
| Cas13a-12a amplification | Cas13a-Cas12a | magnetic bead (MB) tethered with blocker strand (BS) | ~0.35 fM | microRNA (miRNA-155) in clinical cancer samples | 75 min | (3) |
| tanCRISPR | Cas13a-Cas12a | "Locked RNA/DNA" mediator | pM level | microRNA (miRNA-21) in breast cancer cell extracts | ~60 min | (4) |
| An aM-level sensitive cascade CRISPR-Dx system (ASCas) | Cas13a-Cas12a | hybridized cascade probe with RNA bubble | ~1 aM | SARS-CoV-2 RNA extract from pseudovirus samples | ~20 min | (5) |
| CLEAR | Cas13a-Cas12a | Hairpin cascade probe (self-folding) | ~1 aM | SARS-CoV-2 in clinical nasopharyngeal swabs | ~30 min | (6) |
| Hairpin-locker (H-locker) | Cas12a-Cas13a | Asymmetric hairpin "H-locker" | ~1 aM (<1 copy/μL) | DNA (PIK3CA-H1047R mutation) in Plasma (CRC mice) | ~120 min | (7) |
| STACS | Cas13a-Cas12a | linear DNA-RNA-DNA mediator | ~1 copy / μL (~1.67 aM) | certified reference SARS-CoV-2 RNA standards in complex biological and environmental matrices, including serum, saliva, and mud | ~15 min | Our manuscript |
